# FOXA2 controls the anti-oxidant response in FH-deficient cells

**DOI:** 10.1101/2022.07.04.498412

**Authors:** Connor Rogerson, Marco Sciacovelli, Lucas A Maddalena, Lorea Valcarcel-Jimenez, Christina Schmidt, Ming Yang, Elena Ivanova, Joshua Kent, Ariane Mora, Danya Cheeseman, Jason S Carroll, Gavin Kelsey, Christian Frezza

## Abstract

Hereditary Leiomyomatosis and renal cell cancer (HLRCC) is a cancer syndrome caused by inactivating germline mutations in fumarate hydratase (FH) and subsequent accumulation of fumarate. Fumarate accumulation leads to the activation of an anti-oxidant response via nuclear translocation of the transcription factor NRF2. The activation of the anti-oxidant response is key for cellular survival in FH-deficient cells, yet the extent to which chromatin remodelling shapes the anti-oxidant response is currently unknown. Here, we explored the global effects of FH loss on the chromatin landscape to identify transcription factor networks involved in the highly remodelled chromatin landscape of FH-deficient cells. We identify FOXA2 as a key transcription factor which directly regulates anti-oxidant response genes and subsequent metabolic rewiring. Moreover, we also find that FOXA2 regulates anti-oxidant genes independent of the canonical anti-oxidant regulator NRF2. The identification of FOXA2 as an anti-oxidant regulator provides new insights into the molecular mechanisms behind cell responses to fumarate accumulation, and potentially provides new avenues for therapeutic intervention for HLRCC.

## Introduction

Fumarate hydratase (FH) catalyses the reversible conversion of fumarate to malate in the tricarboxylic acid (TCA) cycle. Inactivating germline mutations in *FH* have been identified as the main cause of hereditary leiomyomatosis and renal cell cancer (HLRCC) (Tomlinson et al., 2002), a cancer predisposition syndrome which results in benign skin and uterine lesions and an aggressive form of type II papillary renal cell cancer after loss of heterozygosity. These tumours metastasise early and are poorly survived (Grubb et al., 2007; Menko et al., 2014).

The loss of FH results in a complex rewiring of metabolism, and the accumulation of intracellular fumarate (Frezza et al., 2011), which has been previously linked to the oncogenic potential of FH loss (reviewed in Schmidt et al., 2020). Many of the effects of elevated fumarate are linked to chromatin regulation. Indeed, high levels of intracellular fumarate can competitively inhibit α-ketoglutarate dioxygenases (Laukka et al., 2016; Xiao et al., 2012), leading to a DNA and histone hypermethylation phenotype. DNA hypermethylation is particularly evident in FH-deficient tumours (Sun et al., 2021; The Cancer Genome Atlas Research Network, 2016) and fumarate-induced-DNA methylation at the *mir-200c and mir-200ba429* loci promotes epithelial-to-mesenchymal transition (EMT) in FH-deficient cells by activating the repressor ZEB1 (Sciacovelli et al., 2016). Fumarate accumulation is also able to covalently react with cysteine residues of proteins in a process called succination (Alderson et al., 2006). One of the most well-known targets of succination is KEAP1 (Adam et al., 2011; Ooi et al., 2011), a U3-ubiquitin ligase which targets NRF2 (*NFE2L2* gene codes for NRF2) for proteasomal degradation by ubiquitination (Baird and Yamamoto, 2020). Once KEAP1 is succinated, it releases NRF2 to activate anti-oxidant response genes – another key marker for FH-deficient tumours (Adam et al., 2011; Ooi et al., 2011). As FH-deficient cells exists in a persistent oxidative stress environment due to high levels of fumarate, the activation and fine-tuning of the anti-oxidant response is important for survival (Zheng et al., 2015). This is exemplified by the synthetic lethality of *Hmox1*, a key NRF2 target gene, in FH-deficient cells (Frezza et al., 2011). While the effects of FH loss have often been studied separately, these processes likely occur simultaneously and influence each other. Yet, it is still unclear to what extent the broad chromatin remodelling caused by fumarate synergises with the activation of the above-described transcriptional events.

Chromatin accessibility profiling is well-suited to reflect active signalling pathways that converge on chromatin (Yan et al., 2020). Additionally, H3K27 acetylation marks active promoters and enhancers genome-wide (Creyghton et al., 2010), therefore we can obtain a comprehensive perspective of chromatin regulation by profiling these two features. To ascertain the changes to the chromatin landscape in FH-deficient cells we performed ATAC-seq and H3K27ac ChIP-seq to profile the chromatin landscape of FH-proficient and FH-deficient cells. By integrating these data, we uncovered the key chromatin alterations associated with FH loss and identified important transcription factors in these processes. Unexpectedly, we implicate FOXA2 in the additional regulation of the anti-oxidant response, independent of NRF2. FOXA2 knockdown reduces expression of NRF2 target genes and altered anti-oxidant related metabolites.

## Results

### Dynamic changes in the chromatin landscape of FH-deficient cells

To uncover the regulatory chromatin landscape of FH-deficient cells we performed ATAC-seq and H3K27ac ChIP-seq in *Fh1*-proficient (*Fh1*^*fl/fl*^), two *Fh1*-deficient clones (*Fh1*^*-/-CL1*^ and *Fh1*^*-/-CL19*^) and one *Fh1* rescue cell line (*Fh1*^*-/-CL1*^*+pFh1-GFP*). The ATAC-seq data showed regular nucleosome periodicity and was deemed good quality (Supplementary Figure 1A). The ATAC-seq replicates also correlated well (Supplementary Supplementary Figure 2), as did the H3K27ac ChIP-seq replicates (Supplementary Supplementary Figure 3). We first called peaks on each replicate and merged the peaks together to create two independent union-peak sets for the ATAC-seq dataset (n=112,031) and the H3K27ac ChIP-seq dataset (n=32,161). We performed differential accessibility analysis between *Fh1*^*-/-CL1*^ and *Fh1*^*fl/fl*^, which identified 12,605 differentially accessible regions (Figure 1A; Supplementary Table 1). 7,614 regions (60%) were significantly reduced in *Fh1*^*-/-CL1*^, whereas 4,991 (40%) regions were significantly increased in *Fh1*^*-/-CL1*^ (Figure 1A). Differential H3K27ac ChIP-seq analysis resulted in 12,256 differential regions, of which 6,759 (55%) regions were significantly reduced and 5497 (45%) regions were significantly increased in *Fh1*^*-/-CL1*^ compared to *Fh1*^*fl/fl*^ (Figure 1B; Supplementary Table 2). First, we checked for known oncogenic events in FH-deficiency, and indeed, Vimentin, a marker of EMT, shows an increase in chromatin accessibility and H3K27ac at promoter and distal enhancer regions (Figure 1C), whereas *Mir-200c* shows a marked decrease in H3K27ac and chromatin accessibility at its locus (Supplementary Figure 1D), which is consistent with repression. Additionally, *Slc7a11*, a common target gene of the anti-oxidant response in FH-deficient cells, also shows an increase in H3K27ac and chromatin accessibility in FH-deficient cells at promoter and enhancer regions (Supplementary Figure 1D). These select loci demonstrate that the chromatin landscape is reflecting known biology of FH deficiency. We next examined *Fh1*^*-/- CL19*^ cells, which also show a similar pattern to *Fh1*^*-/-CL1*^ cells, highlighting the ATAC-seq and H3K27ac ChIP-seq data is consistent across clones (Supplementary Figure 1E). To assess whether these chromatin changes were reversible, we profiled the chromatin accessibility and H3K27ac landscape in *Fh1*^*-/-CL1*^ cells stably transfected with a Fh1 expressing plasmid (*Fh1*^*-/-CL1*^*+pFh-GFP*). These cells showed an almost complete rescue of chromatin accessibility and H3K27ac levels (Figure 1D) and clustered more closely to *Fh1*^*fl/fl*^ cells (Supplementary Figure 1B, Supplementary Figure 1C). These results clearly demonstrate a dynamic chromatin reprogramming upon FH loss, which can be reversed with the re-expression of FH.

**Figure 1.**
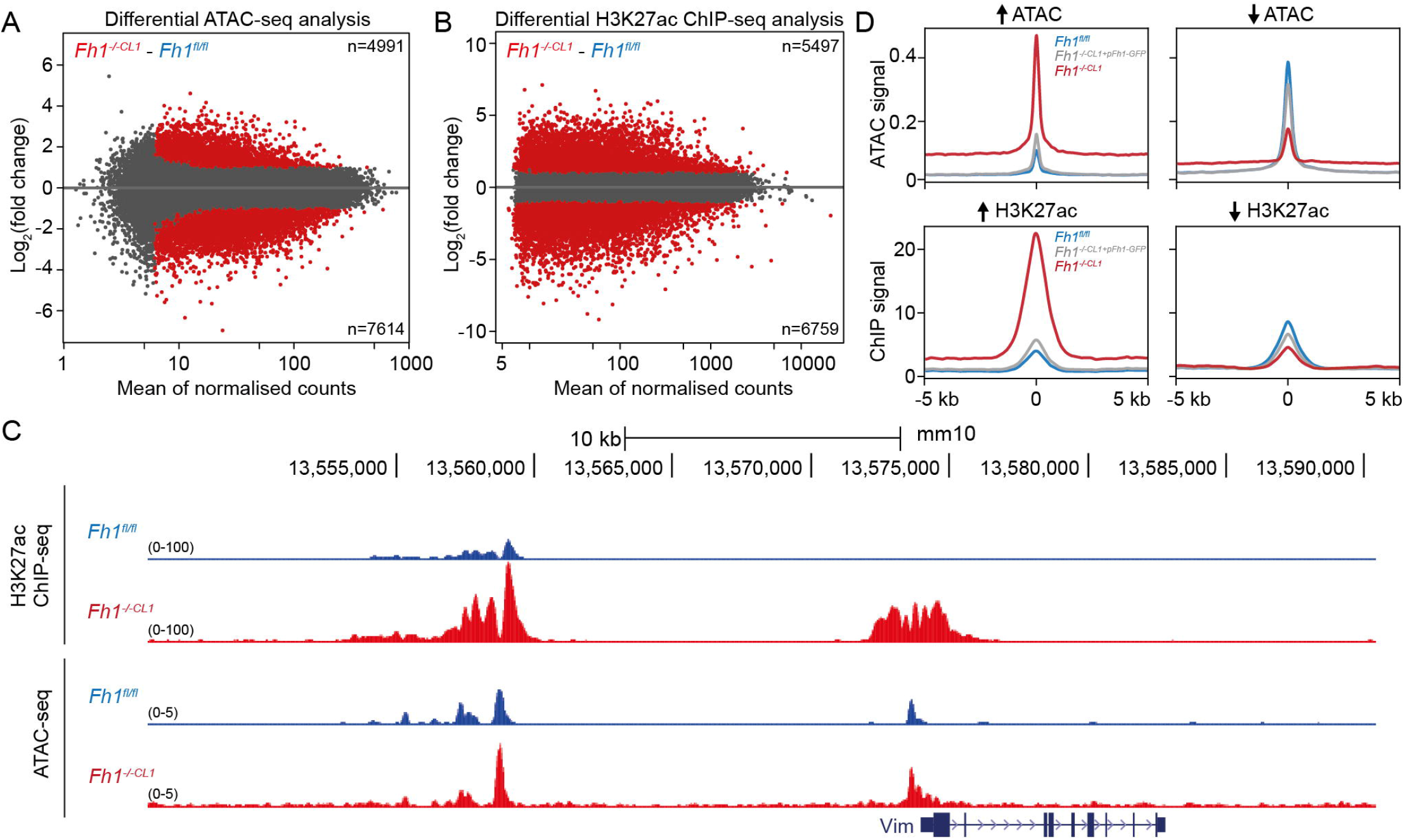
FH-deficiency coincides with chromatin rewiring (A) MA plot of differential ATAC-seq analysis in *Fh1*^*-/-CL1*^ cells compared to *Fh1*^*fl/fl*^ cells. Regions with a linear fold change of ± 2 and q-value of less than 0.05 are highlighted in red. (B) MA plot of differential H3K27ac ChIP-seq analysis in *Fh1*^*-/-CL1*^ cells compared to *Fh1*^*fl/fl*^ cells. Regions with a linear fold change of ± 2 and *q*-value of less than 0.05 are highlighted in red. (C) H3K27ac ChIP-seq and ATAC-seq genome browser tracks from *Fh1*^*fl/fl*^ and *Fh1*^*-/- CL1*^ cells around the *Vim* locus. (D) Normalised ATAC-seq signal (upper) and H3K27ac ChIP-seq signal (lower) from *Fh1*^*fl/fl*^ (blue), *Fh1*^*-/-CL1*^ (red), and *Fh1*^*-/-CL1*^*+pFh1-GFP* (grey) at significantly increased (left) or decreased regions (right).

### Chromatin changes occur in distinct clusters

To gain a further understanding into the mechanisms of gene regulation in FH-deficient cells, we integrated our ATAC-seq and H3K27ac ChIP-seq datasets using simple logical rules. We used the ATAC-seq union-peak set as our reference and assigned to each accessible region whether it first, demonstrated an increase, decrease or no change in chromatin accessibility (±2 fold and *q*-value < 0.05) and second, whether it demonstrated an increase, decrease or no change (±2 fold and *q*-value < 0.05) in H3K27ac ChIP-seq signal (Figure 2A). This resulted in 7 main clusters reflecting the modes of transcriptional regulation (Supplementary Table 3). Open activated regions (I) with an increase in chromatin accessibility and H3K27ac; activated regions (II) with an increase in H3K27ac only; open (III) have an increase in chromatin accessibility only; stable regions (VI) show no change in either chromatin accessibility or H3K27ac; closed regions (V) with a decrease in chromatin accessibility only; deactivated regions (VI) with a decrease in H3K27ac only; and closed deactivated regions (VII) with a decrease in both chromatin accessibility and H3K27ac. A number of regions showed an apparent contradictory pattern: nine regions showed an increase in chromatin accessibility and a decrease in H3K27ac; and 26 regions showed a decrease in chromatin accessibility and an increase in H3K27ac. As these regions represented less than 0.0003 % of the total dataset, they were excluded from further analyses.

**Figure 2.**
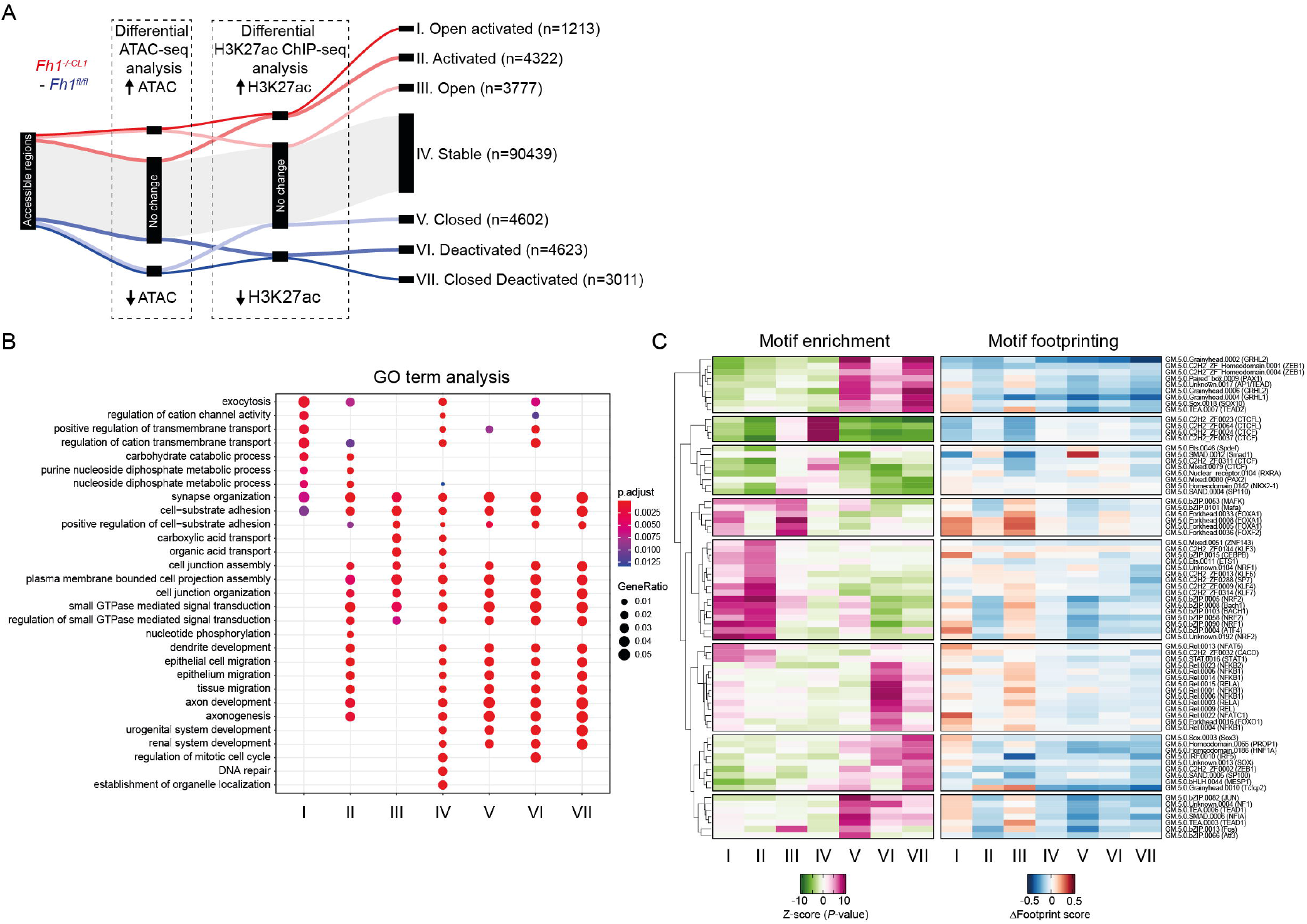
Integrating ATAC-seq and H3K27ac data identify distinct clusters of regulation (A) Sankey diagram of the process of classifying each accessible region in clusters depending on an increase, decrease or no change in ATAC-seq signal and an increase, decrease or no change in H3K27ac ChIP-seq signal. (B) Dot plot of enriched gene ontology (GO) terms of genes located near to regions of clusters I – VII. Size of dots represents proportion of differentially expressed genes in a given GO term (GeneRatio) and colours represents significance (p.adjust) (C) Heatmap of enrichment of transcription factor motifs (left) and differential footprinting score (right) within regions I – VII.

The increase in ATAC-seq signal was evident in clusters I and III, whereas a decrease in ATAC-seq signal was clear in clusters V and VII (Supplementary Figure 2A). Similarly, H3K27ac ChIP-seq increases can be seen in clusters I and II, and decreases in clusters VI and VII (Supplementary Figure 2B). The distance distribution to the nearest transcription start site (TSS) was similar for all clusters and the majority of regions were distal to a TSS (Supplementary Figure 2C). Some clusters showed a large proportion of regions falling within 3 kb from a TSS, chiefly cluster II (activated) regions. To see whether these changes in chromatin accessibility and activation translated into transcriptional changes we interrogated previously published RNA-seq datasets from FH-deficient cells (Sciacovelli et al., 2016). Importantly, the regions within these clusters are significantly associated with transcriptional changes in FH-deficient cells, compared to a random geneset (Supplementary Figure 2D). More active regions (clusters I, II and III) are significantly associated with upregulated nearby genes, and less-active regions (clusters V, VI and VII) are significantly associated with downregulated nearby genes (Supplementary Figure 2), showing that the logical rules we set were able to group genes into meaningful clusters. Moreover, the clusters demonstrate that underpinning the dynamic chromatin landscape are discrete modes of regulation involving chromatin accessibility and histone modifications that are linked to gene expression.

### Transcription factor motifs define clusters

To assess whether these clusters of regions were associated with distinct sets of genes, we performed GO (Gene Ontology) term analysis of the nearest genes associated with each region and cluster (Figure 2B). Known activated processes involved in FH-deficiency include EMT, anti-oxidant stress signalling and amino acid metabolism. Cluster I regions (open activated) were enriched in GO terms involving nucleoside, carbohydrate metabolic processes and transmembrane transport. These GO terms are important processes in amino acid metabolism and anti-oxidant response in general. Cluster II regions (activated) were enriched in migration and cell adhesion related GO terms, which are heavily associated with EMT. Cluster III regions (open) were also enriched in metabolite transport, migration and GTPase signal transduction GO terms.

Cluster V, VI and VII regions (closed, deactivated and closed deactivated) were enriched in renal development and cell cycle GO terms. Loss of kidney-specific developmental genes is perhaps reflecting a dedifferentiation process common in tumours (Friedmann-Morvinski and Verma, 2014). Furthermore, the presence of cell cycle GO term enrichment in repressed clusters is consistent with previous observations that FH-deficient cells have a proliferative defect (Frezza et al., 2011).

The specific GO terms associated with the clusters indicate that these regions are regulating genes involved in specific functions and pathways known in FH deficiency. To link putative transcription factors to the regulation of these genes, we performed *de novo* motif analysis on each cluster (Figure 2C; Supplementary Table 4). Strikingly, this revealed very specific enrichment of transcription factors for each cluster. Cluster IV regions (stable) were highly enriched for CTCF motifs, suggesting that these regions represent stable structural elements of the genome. Cluster I and II regions (open activated/activated) were enriched for NRF2 and ATF4 motifs, transcription factors known to induce the anti-oxidant response, whereas cluster III regions (open) were highly enriched in forkhead motifs. In fact, forkhead motifs were enriched in both cluster I and II regions, suggestive that forkhead motifs are perhaps important for the activity of cluster I transcription factor motifs i.e. NRF2 or ATF4. Cluster V regions (closed) were enriched in AP-1 and TEAD motifs. Cluster VI regions (deactivated) were enriched in NFKB motifs and cluster VII regions (closed deactivated) were enriched in grainyhead-like and ZEB motifs. Grainyhead transcription factors (e.g. GRHL2) specify the kidney and maintain its identity (Boivin and Schmidt-Ott, 2020), whereas ZEB is a repressor involved in repressing epithelial genes and contributing to EMT (Zhang et al., 2015) – this is consistent with the renal development and migratory GO terms enriched in clusters V, VI and VII. We also took advantage of the ATAC-seq data to analyse footprinting around transcription factor motifs (a proxy for transcription factor binding). Our analysis shows a general increase in footprinting in cluster I and III, especially at forkhead motifs, but also ATF4 and NRF2 motifs (Figure 2C; Supplementary Table 4). Conversely, Grainyhead-like motifs showed the largest reduction in footprinting. Together, these data link the regulation of genes at the chromatin level to presence of specific transcription factors and their motifs, and points towards a complex network of transcription factors that underly the response to FH loss.

### FOXA2 regulates NRF2 target genes in FH-deficient cells

We were interested in additional novel transcription factors that respond to FH loss, therefore we focussed on a putative forkhead transcription factor that may be acting in concert with other transcription factors. We first looked at the level of expression of forkhead factors *Fh1*^*-/-CL1*^ cells that showed a significant change in expression compared to *Fh1*^*fl/fl*^ cells (Supplementary Figure S3A), however there was not clear highly expressed transcription factor. Additionally closer inspection of the magnitude of the fold change in gene expression changes did not identify a clear single transcription factor (Supplementary Figure S3B). We then mined previously published CRISPR-screening data (Valcarcel-Jimenez et al., 2022) – here a genome-wide pooled CRISPR screen was performed on *Fh1*^*fl/fl*^ and *Fh1*^*-/-CL1*^ cells to identify essential genes in cell survival. The two screens were compared to each other to identify essential genes specifically in FH-deficient cells. Using this data we were able to identify *Foxa2* as a key gene important in survival only in *Fh1*^*-/-CL1*^cells (Figure 3A). *Foxa2* is moderately expressed in *Fh1*^*fl/fl*^ cells and its expression is increased in *Fh1*^*-/-CL1*^ cells at the mRNA (Figure 3B) and protein level (Figure 3C). Additionally *Foxa2* mRNA expression is increased on the protein level (Figure 3C). Importantly, the expression of *Foxa2* in *Fh1*^*-/- CL1*^*+pFh-GFP* cells is completely abrogated, highlighting the reversibility of *Foxa2* expression with the re-expression of FH.

**Figure 3.**
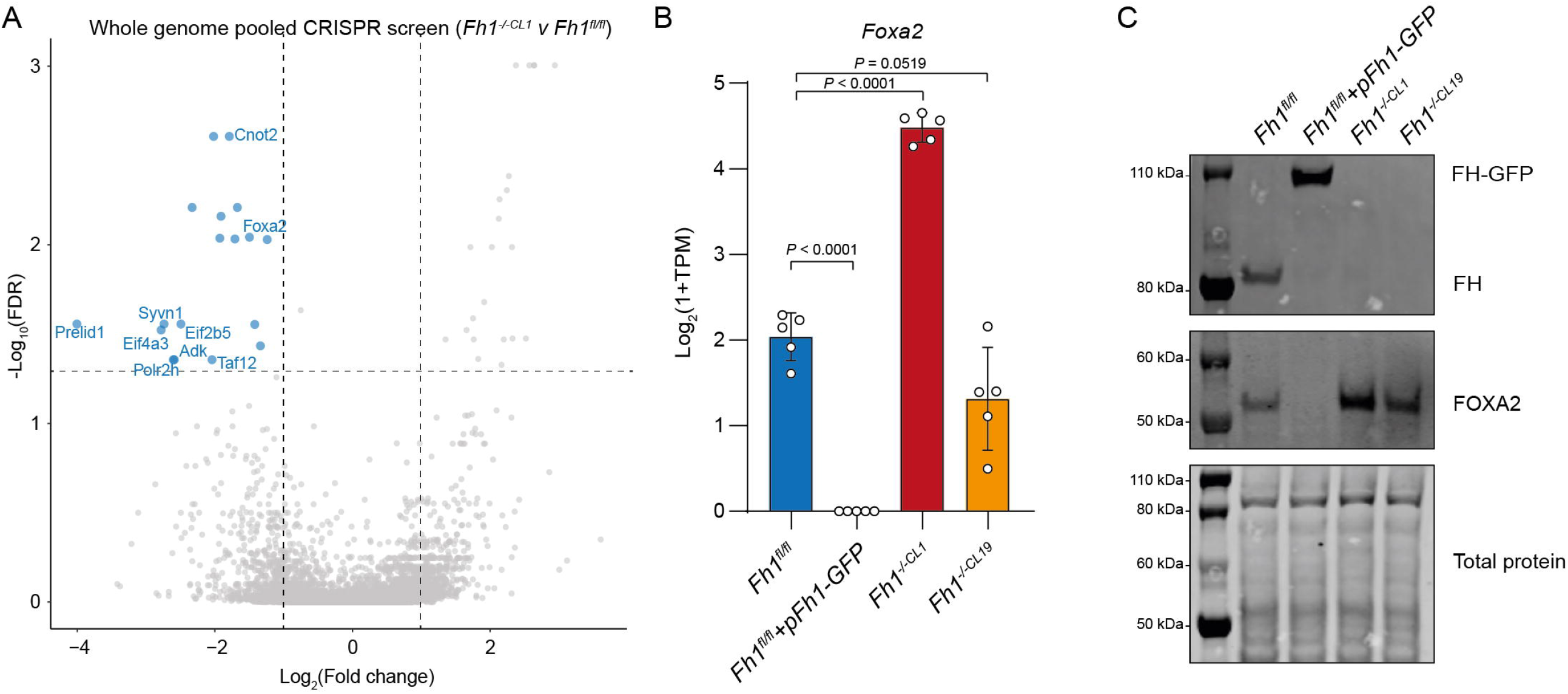
Whole genome CRISPR screen identifies FOXA2 as an important transcription factor (A) Volcano plot of genes enriched in a whole genome CRISPR screen in *Fh1*^*fl/fl*^ and *Fh1*^*-/-CL1*^ cells. The comparison was made between *Fh1*^*fl/fl*^ and *Fh1*^*-/-CL1*^ cells to identify lethal genes dependent on FH-deficiency. Points highlighted in blue are significantly depleted hits and labelled points are high confidence hits (defined as enriched compared to control cells taking into account original plasmid). (B) Expression of *Foxa2* in *Fh1*^*fl/fl*^ (blue), *Fh1*^*-/-CL1*^*+pFh1-GFP* (grey), *Fh1*^*-/-CL1*^ (red) and *Fh1*^*-/-CL19*^ (orange). (C) Immunoblot of protein lysate from *Fh1*^*fl/fl*^, *Fh1*^*-/-CL1*^*+pFh1-GFP, Fh1*^*-/-CL1*^ and *Fh1*^*-/-CL19*^ cells probed with antibodies against FOXA2 and FH.

We further examined the expression *FOXA2* in HLRCC patient samples. *Foxa2* was overexpressed in FH-deficient human tumours, compared to adjacent normal tissue (Supplementary Figure 3C; Crooks et al., 2021). We then examined TCGA data from the papillary renal cell carcinoma dataset (KIRP), the histological subtype of HLRCC, and compared FH negative and FH positive tumours (see methods). Importantly, *Foxa2* expression was increased in FH negative tumours, underscoring the increase in Foxa2 expression is seemingly dependent on FH loss (Supplementary Figure 3D). Overexpression of Foxa2 is also associated with worse overall survival in TCGA KIRP patients (Supplementary Figure 3E).

To ascertain the role of FOXA2, we performed siRNA-mediated silencing of *Foxa2* in *Fh1*^*fl/fl*^, *Fh1*^*-/-CL1*^ and *Fh1*^*-/-CL19*^ cells (Supplementary Figure 4A; Supplementary Table 5) and performed RNA-seq, The RNA-seq replicates clustered well with each other (Supplementary Figure 4B). Differential gene expression analysis between siFoxa2 and siNT in *Fh1*^*fl/fl*^, *Fh1*^*-/- CL1*^ and *Fh1*^*-/-CL19*^ cells identified 119, 838 and 545 differentially expressed genes (DEGs) respectively (Figure 4A). This result highlights the specificity of FOXA2 in FH-deficient cells. Importantly, 57% (311/545; *P* < 7.177e-263) of *Fh1*^*-/-CL19*^ siFoxa2 DEGs were also DEGs in siFoxa2 treated *Fh1*^*-/-CL1*^ cells (Supplementary Figure 4C), underscoring that the function of FOXA2 is similar in both FH-deficient cell-lines.

**Figure 4.**
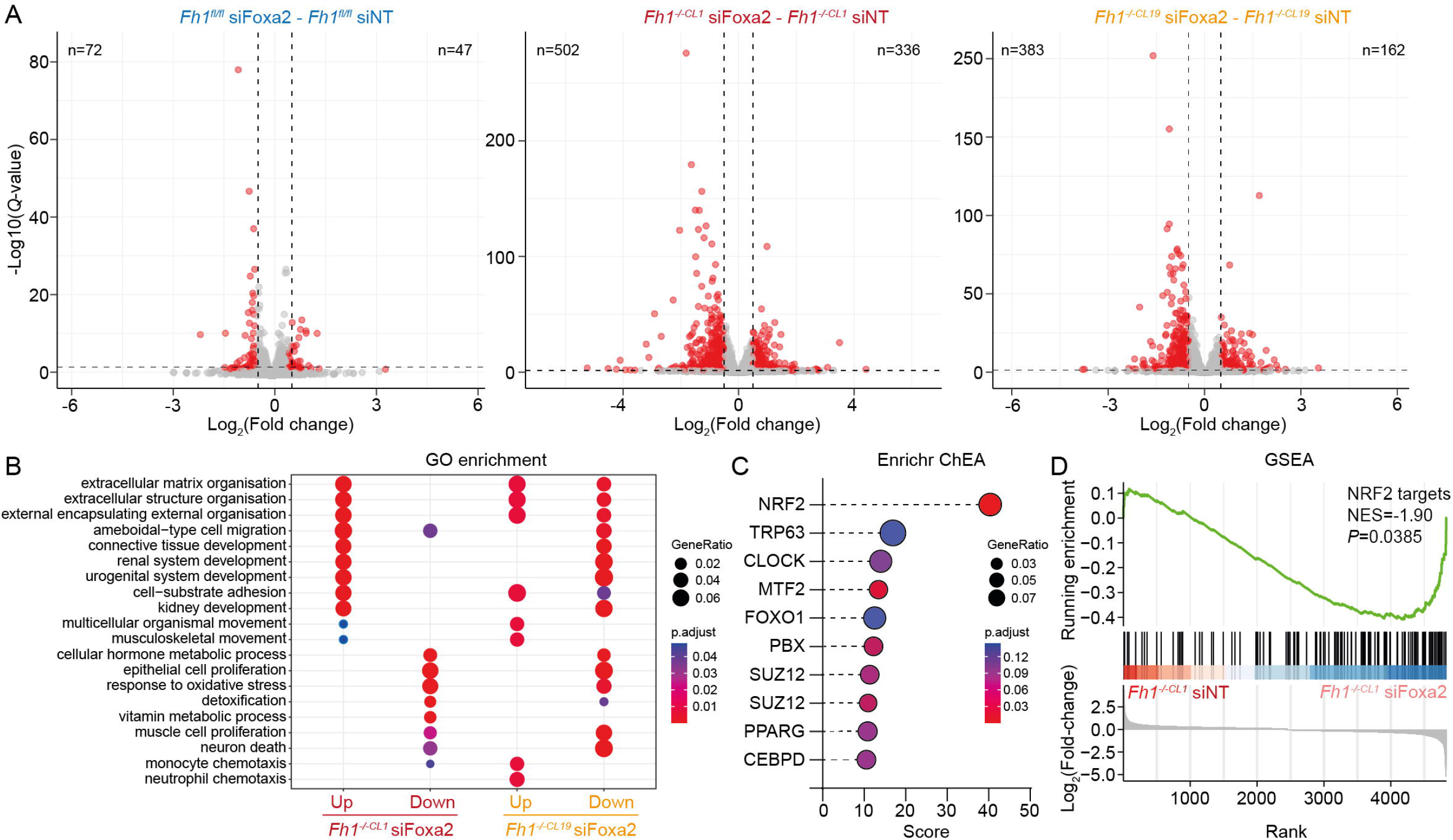
FOXA2 controls NRF2 target genes (A) Volcano plots of differentially expressed genes in *Fh1*^*fl/fl*^ (left), *Fh1*^*-/-CL1*^ (middle) and *Fh1*^*-/- CL19*^ (right) cells treated with siFoxa2 compared to siNT. (B) Gene ontology enrichment of differentially expressed genes in *Fh1*^*-/-CL1*^ (left) and *Fh1*^*-/-CL19*^ (right) cells. (C) Transcription factor enrichment (ChEA) from publicly available ChIP-seq datasets nearby differentially expressed genes in *Fh1*^*-/-CL1*^ cells treated with siFoxa2 compared to siNT. (D) Enrichment of NRF2 target genes (NFE2L2.V2 gene set) among differentially expressed target genes in *Fh1*^*-/-CL1*^ cells treated with siFoxa2 compared to siNT.

To understand what process FOXA2 may regulate, we performed GO term enrichment analysis on differentially regulated genes (Figure 4B). siFoxa2 upregulated genes were enriched in GO terms for extracellular matrix organisation, migration and renal development terms, whereas siFoxa2 downregulated genes were enriched for proliferation, oxidative stress response and detoxification GO terms. Although siFoxa2 upregulated GO terms were not shared between clones (likely indirect effects), siFoxa2 downregulated GO terms were also enriched in *Fh1*^*-/-CL19*^ DEGs (Figure 4B). We then asked whether these DEGs are associated with the original clusters identified from the ATAC-seq and H3K27ac ChIP-seq data. Downregulated genes after siFoxa2 treatment are significantly associated to clusters I and II (open activated/activated), and upregulated genes to clusters VI and VII (deactivated/closed deactivated) (Supplementary Figure 4D), suggesting that FOXA2 directly regulates key processes associated with FH deficiency.

FOXA2 is a pioneer factor in multiple cell types (Zaret and Carroll, 2011). To explore whether FOXA2 may be regulating genes as a pioneer factor for another transcription factor, we used our siFoxa2 downregulated gene list to check whether other transcription factors regulate the same genes. Using ChEA (a database that mines public ChIP-seq datasets), the top enriched transcription factor was NRF2 (Figure 4C), a master redox transcription factor already implicated in FH-deficient cells and tumours (Adam et al., 2011; Ooi et al., 2011). NRF2 target genes were also highly enriched in our siFoxa2 dataset (Figure 4D). These results show that FOXA2 regulates genes involved in active processes in FH-deficient cells, and that FOXA2 is able to regulate NRF2 target genes.

### FOXA2-mediated gene regulation is NRF2 independent

To gain an insight into the mechanism by which FOXA2 regulates NRF2 target genes, we performed FOXA2 ChIP-seq in *Fh1*^*fl/fl*^ and *Fh1*^*-/-CL1*^ cells. The ChIP-seq replicates correlated well and were merged (Supplementary Figure 5A and S5B). Peak calling identified 385 FOXA2 peaks in *Fh1*^*fl/fl*^ cells and FOXA2 4273 peaks in *Fh1*^*-/-CL1*^ cells (Figure 5A; Supplementary Table 6), highlighting its heightened activity in FH-deficient cells. Only 5% (213/4273) of peaks overlapped between the datasets, thus we focussed on *Fh1*^*-/-CL1*^ specific FOXA2 peaks (Figure 5B). Globally, FOXA2 binding regions coincide with an increase in chromatin accessibility and H3K27ac ChIP-seq signal only in *Fh1*^*-/-CL1*^ specific FOXA2 peaks and shared peaks (Supplementary Figure 5C). An example is the *Gclc* locus, which shows increases of H3K27ac ChIP-seq signal and ATAC-seq signal at promoter and upstream distal regions. FOXA2 can be seen to bind specifically in *Fh1*^*-/-CL1*^ cells at the most distal region, accompanied by an increase in ATAC-seq signal (Figure 5C). *Gclc* is a NRF2 target gene (Li et al., 2009), therefore we analysed whether NRF2 target genes (identified from the previous GSEA analysis) were generally bound by FOXA2 in *Fh1*^*-/-CL1*^ cells - FOXA2 ChIP-seq peaks were indeed significantly enriched nearby NRF2 target genes (Supplementary Figure 5D).

**Figure 5.**
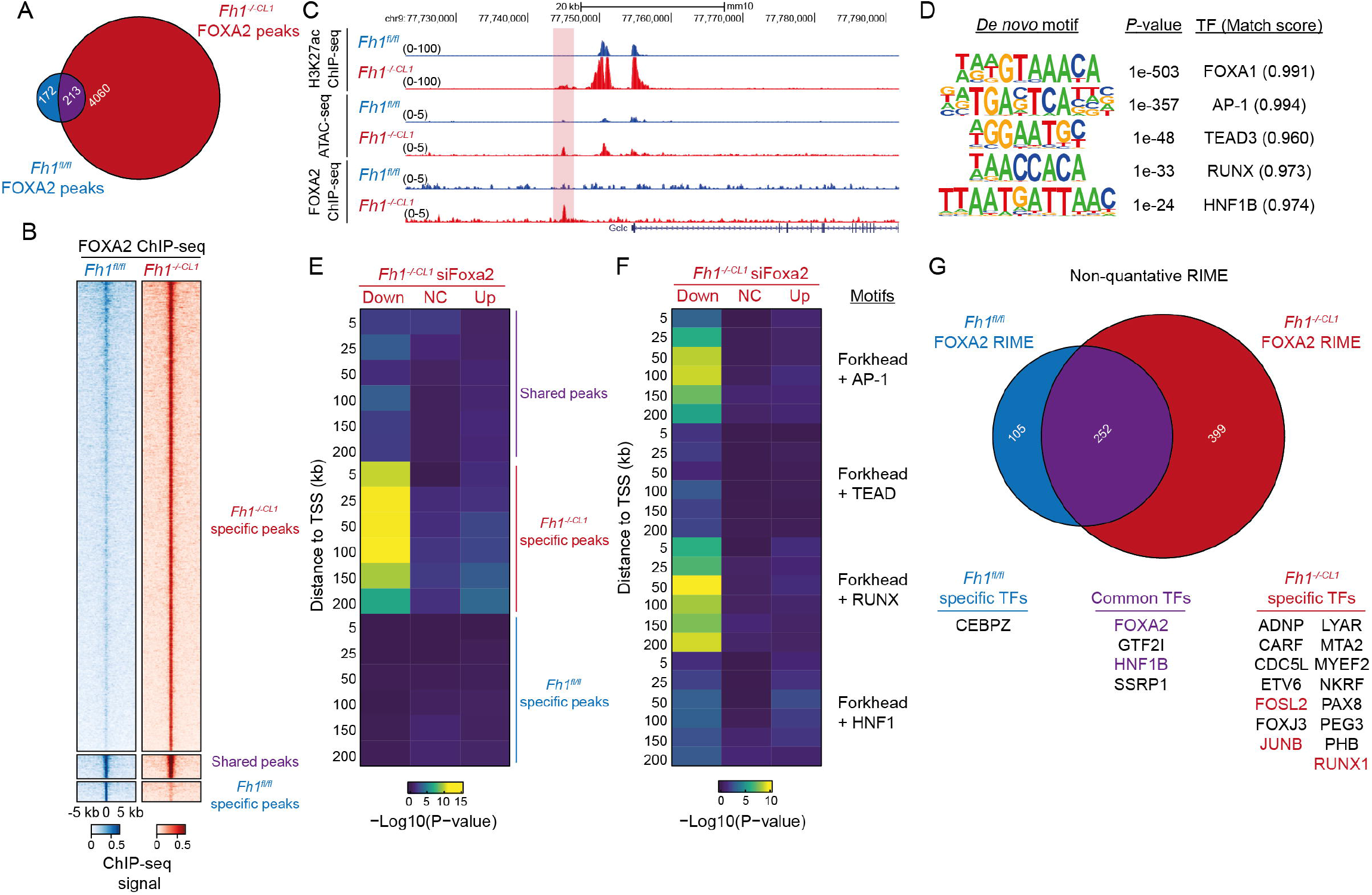
FOXA2 binds chromatin in FH-deficient cells independent of NRF2 (A) Venn diagram of FOXA2 ChIP-seq peaks shared between (purple) and specific to *Fh1*^*fl/fl*^ (blue) and *Fh1*^*-/-CL1*^ cells (red). (B) Heatmap of FOXA2 ChIP-seq signal across 10 kb regions centered at FOXA2 peaks in *Fh1*^*fl/fl*^ (blue) and *Fh1*^*-/-CL1*^ cells (red). (C) H3K27ac ChIP-seq, ATAC-seq and FOXA2 ChIP-seq from *Fh1*^*fl/fl*^ (blue) and *Fh1*^*-/-CL1*^ cells (red) at the *Gclc* locus. *Fh1*^*-/-CL1*^ specific FOXA2 peak upstream of *Gclc* highlighted in red. (D) Enriched motifs from *de novo* motif analysis of FOXA2 peaks specific for *Fh1*^*-/-CL1*^ cells with enrichment *P*-values and matched transcription factor names and match scores. (E) Heatmap of enrichment *P*-values (hypergeometric test) of differentially expressed genes from siFoxa2 treatment to FOXA2 ChIP-seq peaks at increasing distances from the TSS. (F) Heatmap of enrichment *P*-values (hypergeometric test) of differentially expressed genes from siFoxa2 treatment to FOXA2 ChIP-seq peaks with the presence additional transcription factor motifs at increasing distances from the TSS. (G) Venn diagram comparing proteins identified from FOXA2 RIME in *Fh1*^*fl/fl*^ (blue) and *Fh1*^*-/-CL1*^ cells (red). Proteins from each section that are annotated as a transcription factor are stated, and highlighted if their motif was identified from the *de novo* motif analysis.

To gain an insight into the mechanism of gene activation by FOXA2, we performed *de novo* motif analysis on *Fh1*^*-/-CL1*^ specific FOXA2 peaks to identify additional enriched. Reassuringly the top enriched motif was the forkhead motif, highlighting that the target was correctly pulled down. The other enriched motifs were for AP-1, TEAD, RUNX and HNF1 (Figure 5D). As the consensus motif for NRF2 was not identified we profiled the occurrence of the NRF2 motif within *Fh1*^*-/-CL1*^ specific FOXA2 peaks compared with forkhead, AP-1 and CTCF motifs. As expected, these peaks were highly enriched in motifs for AP-1 and FOXA2, but surprisingly completely devoid of NRF2 motifs (Supplementary Figure 5D). We next assessed the connection of *Fh1*^*-/-CL1*^ specific FOXA2 peaks to target genes identified from the siFoxa2 RNA-seq data. FOXA2 ChIP-seq peaks were significantly enriched nearby to siFoxa2 downregulated genes (Figure 5E), and these regions were likely to house AP-1 and/or RUNX motifs in addition to a forkhead motif (Figure 5F).

While this analysis did not identify the NRF2 motifs, FOXA2 could potentially still tether NRF2 to its target. To test this hypothesis, we performed rapid immunoprecipitation mass spectrometry of endogenous protein (RIME) of FOXA2 in *Fh1*^*fl/fl*^, *Fh1*^*-/-CL1*^ cells to identify interactors of FOXA2 on chromatin (Supplementary Table 7). All RIME replicates identified FOXA2 highlighting the validity of the datasets, and proteins found in at least 2 or more replicates were taken forward. This resulted in 105 *Fh1*^*fl/fl*^ specific interactors, 252 shared interactors and 399 *Fh1*^*-/-CL1*^ specific interactors (Figure 5G). We further filtered these proteins for transcription factors and identified CEPBZ in *Fh1*^*fl/fl*^ only, FOXA2 and HNF1B shared between both *Fh1*^*-/-*^ cell-lines, and AP-1 (FOSL2, JUNB) and RUNX1 only in *Fh1*^*-/-CL1*^ cells. Still, we did not identify NRF2 as an interactor in this dataset. Together these data suggest that FOXA2 can regulate NRF2 target genes independent of NRF2 itself.

### FOXA2 regulates NRF2 related metabolic pathways

Thus far the data suggest that FOXA2 converges on similar genes and pathways as NRF2, yet the mechanism of action is independent of NRF2. Finally, to assess whether there is a functional convergence between the two proteins, we performed metabolomics on *Fh1*^*fl/fl*^, *Fh1*^*-/-CL1*^ and *Fh1*^*-/-CL19*^ cells treated with either siNT or siFoxa2 (Supplementary Table 7). The knockdown was successful (Figure 6A) and metabolomic replicates clustered together (Supplementary Figure 6A). Performing differential metabolite analysis, we identified various metabolites whose abundances were significantly altered in siFoxa2 treated cells compared to siNT (Figure 6B). *Fh1*^*-/-*^ cell lines demonstrated a deeper metabolic reprogramming than *Fh1*^*fl/fl*^ cells upon *Foxa2* silencing, consistent with FOXA2 being more active in FH-deficient cells.

**Figure 6.**
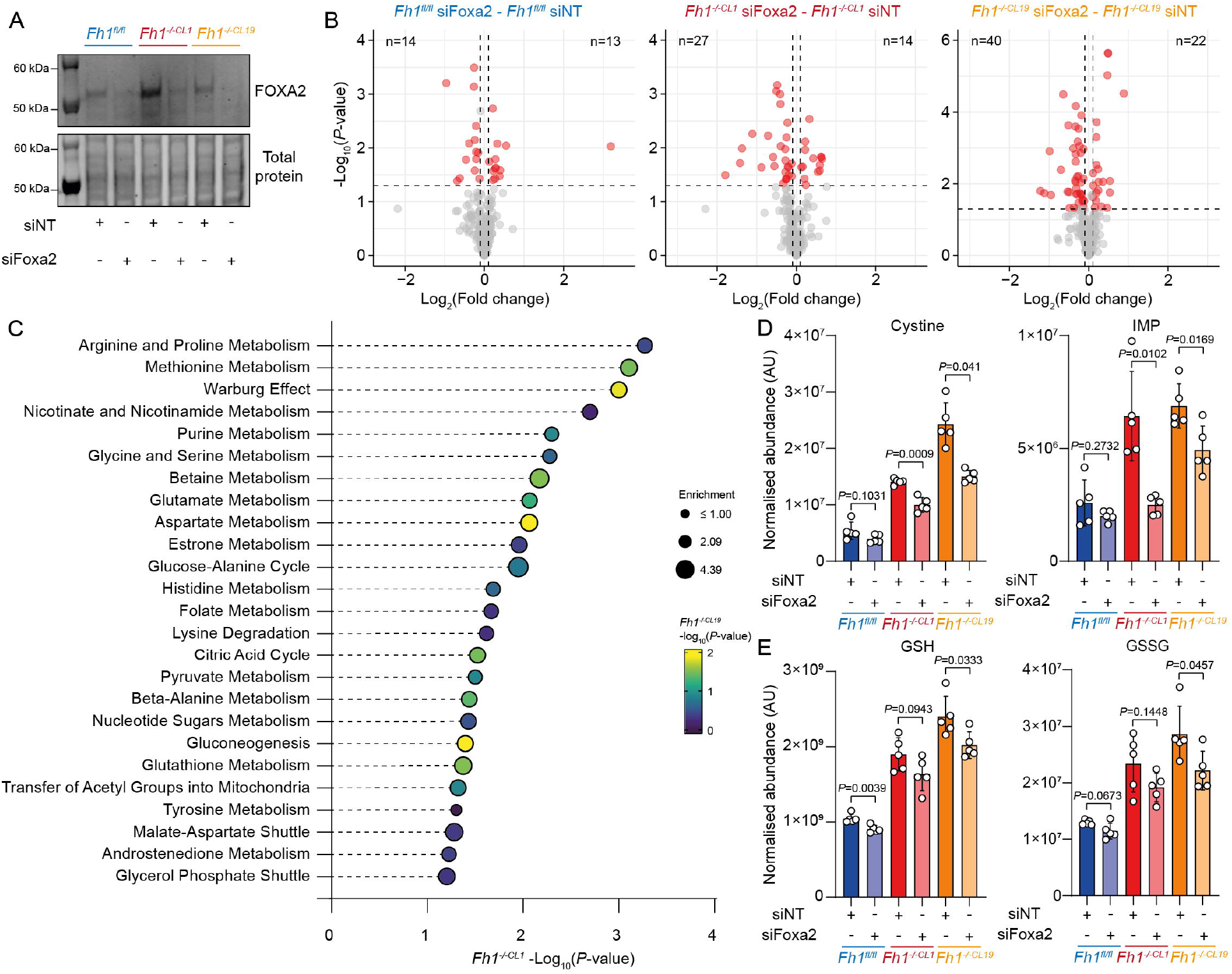
FOXA2 regulates NRF2 associated metabolism (A) Immunoblot of protein lysate from *Fh1*^*fl/fl*^, *Fh1*^*-/-CL1*^ and *Fh1*^*-/-CL19*^ cells treated with either siNT or siFoxa2 probed with antibodies against FOXA2. (B) Volcano plots of differentially abundant metabolites in *Fh1*^*fl/fl*^ (left), *Fh1*^*-/-CL1*^ (middle) and *Fh1*^*-/-CL19*^ (right) cells treated with siFoxa2 compared to siNT. (C) Metabolite set enrichment analysis of significantly changed metabolites in *Fh1*^*-/-CL1*^ cells treated with siFoxa2 compared to siNT. Colour of dots represent enrichment in altered metabolites in *Fh1*^*-/-CL19*^ cells treated with siFoxa2 compared to siNT. (D-E) Normalised abundances of indicated metabolites in *Fh1*^*fl/fl*^, *Fh1*^*-/-CL1*^ and *Fh1*^*-/-CL19*^ cells treated with either siNT or siFoxa2.

To identify metabolic pathways that were altered in FH-deficient cells and dependent on FOXA2, we used metabolite set enrichment analysis (Figure 6C). This identified numerous pathways, including methionine, glutamate and aspartate metabolism, gluconeogenesis and glutathione metabolism. Importantly, these pathways were also enriched in *Fh1*^*-/-CL19*^ cells. We then focussed on individual metabolites known to be regulated by NRF2 (DeBlasi and DeNicola, 2020). Indeed, cystine and inositol monophosphate (IMP) were significantly reduced specifically in siFoxa2 treated *Fh1*^*-/-*^ cells, whilst in siFoxa2 treated *Fh1*^*fl/fl*^ cells they were not reduced (Figure 6D). The levels of glutathione (GSH) and glutathione-disulfide (GSSG) were also reduced in siFoxa2 treated cells, whilst the GSH/GSSG ratio did not change (Figure 6E).

Collectively, these results show that FOXA2 can functionally regulate the output of select metabolic pathways associated with NRF2 activation.

## Discussion

The consequences of FH loss are still poorly understood after it was identified as the cause of HLRCC two decades ago (Tomlinson et al., 2002). Recent studies clearly point to an intimate relationship between fumarate and chromatin. Fumarate is able to inhibit DNA demethylases and histone demethylases, thereby increasing methylated DNA and histone levels (Xiao et al., 2012), the majority of which are repressive marks. The importance of increases in DNA methylation is well demonstrated in its role in inducing EMT in FH-deficient cells (Sciacovelli et al., 2016). Fumarate’s ability to inhibit histone demethylases also has a role in DNA repair. Local generation of fumarate by nuclear FH increases histone H3K36 levels to promote DNA repair at double strand breaks (Jiang et al., 2015). Additionally, protein succination from high fumarate levels has been identified on SMARCC1, a key subunit of the SWI/SNF chromatin remodelling complex (Kulkarni et al., 2019). Fumarate is also able to activate anti-oxidant transcription factor NRF2 via KEAP1 succination (Adam et al., 2011) and integrated stress response transcription factor ATF4 (Ryan et al., 2021). A global outlook on regulatory chromatin dynamics in FH-deficient cells has since been overlooked. In this study, we sought to use unbiased approaches to profile the open chromatin landscape (ATAC-seq) and its activity (H3K27ac ChIP-seq). Indeed, we show that the changes in the chromatin landscape are dynamic and widespread: the majority of regions show a decrease in chromatin accessibility (60%) and H3K27ac ChIP-seq (55%); and a substantial number of regions show an increase in chromatin accessibility (40%) and H3K27ac ChIP-seq (45%).

By integrating chromatin accessibility and H3K27ac ChIP-seq using logical rules, we were able to cluster genomic regions by their apparent increase or decrease in chromatin accessibility and activity (H3K27ac). These regions were able to be associated with genes involved in processes known to be active in FH-deficient cells e.g. EMT (Sciacovelli et al., 2016). New associated pathways were also identified, such as renal development. This is consistent with a recent appreciation that cell dedifferentiation and loss of cellular identity factors are a more common feature of human cancers (Friedmann-Morvinski and Verma, 2014). We also took advantage of the underlying DNA sequence of these regions and identified an array of enriched transcription factor motifs. Among the enriched motifs in more active regions were NRF2, ATF4 and MAF motifs. These motifs map to transcription factors known to be active in FH-deficient cells (Adam et al., 2011; Ooi et al., 2011; Ryan et al., 2021). Conversely, ZEB, GRHL and NFKB/RELA motifs were enriched in less active regions. Again, since ZEB1 is involved in repression of epithelial genes during EMT (Thiery et al., 2009), it is consistent that we find ZEB motifs within repressed regions.

We also identified forkhead motifs as enriched in regions associated with opening, consistent with the pioneer factor function of forkhead transcription factors (e.g. FOXA) in other cell types (Zaret and Carroll, 2011). Importantly, we also specifically identified FOXA2 as a hit in a comparative CRISPR screen in FH-deficient cells compared to FH-proficient cells (Valcarel-Jiminez et al., 2022), and its expression is upregulated in FH-deficient cells. FOXA2 expression is also increased in FH-deficient HLRCC tumours, and patients with high FOXA2 expression have an overall poorer prognosis than those with low expression.

FOXA2 is a pioneer factor (Iwafuchi-Doi et al., 2016) and has been associated with metabolic regulation in other cell types (Gao et al., 2010; Wolfrum et al., 2004; Zhang et al., 2005), but never associated with FH loss. Exactly how FH loss results in FOXA2 activation is very interested phenomenon and will warrant further study. Here, we implicated FOXA2 in the regulation of genes involved in cell growth, kidney development and cellular detoxification. More specifically, FOXA2 regulated genes showed substantial overlap with classic NRF2-dependent target genes involved in the anti-oxidant response. Usually, NRF2 regulates its target genes by its inhibitory partner KEAP1 becoming degraded after ROS signalling and then translocating into the nucleus to directly bind anti-oxidant response elements. Our FOXA2 ChIP-seq dataset fails to identify the NRF2 motif as an enriched motif, but does identify the more common AP-1 motif. AP-1 and NRF2 motifs do share the canonical TGA^C^/_G_TCA motif, and NRF2 ChIP-seq datasets have in fact identified the AP-1 motif (Malhotra et al., 2010). In this context, perhaps FOXA2 allows NRF2 to bind less favourable AP-1 sites. However, our RIME data also fails to identify NRF2 as an interactor, but does identify transcription factors that bind enriched motifs in our FOXA2 ChIP-seq data, such as RUNX1, FOSL1, and HNF1B. In this context, FOXA2 may function as a transcription factor on its own accord, rather than a pioneer factor for other transcription factors, and support the anti-oxidant response in concert with canonical anti-oxidant response regulators e.g. NRF2.

FOXA2 can also significantly affect various metabolic pathways. Here we show metabolomics data of cells deficient in FH and FOXA2 revealing impaired amino acid import and metabolism, with glutathione metabolism also being affected. *Slc7a11* codes for a key cysteine transporter, which is heavily activated during the anti-oxidant response as cystine is used as a precursor to the glutathione pathway (Zheng et al., 2015). *Foxa2* knockdown is able to reduce the expression of cystine importer *Slc7a11* and reduce the levels of intracellular cystine, thus impairing the levels of GSH and GSSG overall. The fine-tuning of the anti-oxidant response in FH-deficient cells is paramount to cellular survival, and could be a key step in early transformation (reviewed in Schmidt et al., 2020). With this in mind, it seems likely that key pathways are supported by multiple regulators, and FOXA2 seems to be one of them.

The novel link between FOXA2, NRF2 and the anti-oxidant response provokes questions about whether this is a kidney specific mechanism. In fact, *Foxa2* is induced in *Nrf2*^*-/-*^ lung type II cells when exposed to GSH (Reddy et al., 2007) suggesting a similar mechanism might exist in lung cells, where FOXA2 can induce the anti-oxidant response.

In summary, using a multi-omic approach we have uncovered a role of FOXA2 in regulating the anti-oxidant response, independent of NRF2. We have identified new players in the anti-oxidant response and new avenues for therapeutic exploration in HLRCC. We have also highlighted the importance of unbiased, genome-wide techniques in understanding complex parallel signalling mechanisms.

## Methods

### Cell culture

*Fh1*^*fl/fl*^,*Fh1*^*-/-CL1*^,*Fh1*^*-/-CL19*^ were generated from mouse kidney epithelial cells as described previously (Frezza et al., 2011) and *Fh1*^*-/-CL1*^*+pFh1-GFP* cells were generated by stably transfecting *Fh1*^*-/-CL1*^ cells with a plasmid expression Fh1 as described previously (Sciacovelli et al., 2016). Cells were cultured in DMEM supplemented with 10% FBS. Cells were routinely tested negative for mycoplasma infection and were authenticated by short tandem repeat analysis.

### ATAC-seq

ATAC-seq was carried out as described previously (Corces et al., 2017). 5×10^5^ *Fh1*^*fl/fl*^ cells or 1×10^6^ *Fh1*^*-/-*^ cells were plated onto 6-cm dishes and once reached 80% confluency, detached using trypsin (0.25% in PBS-EDTA). 1×10^4^ cells were then resuspended in cold lysis buffer (10mM TrisCl pH 7.4, 10mM NaCl, 3mM MgCl2, Nonidet P40 0.1%) on ice and centrifuged at 500 rcf/10 min. The cell pellet was then mixed and incubated at 37°C for 1h min with the transposition reaction mix prepared using Illumina Tagment DNA Enzyme and Buffer kits (Illumina, 20034197). DNA was then purified using DNA Clean and Concentrator kit (Zymo, D4013) and library preparation and purification performed using KAPA HiFi Hot Start DNA Polymerase (KAPA Biosystems, KK2500) and Ampure XP beads (Beckman Coulter, A63880). Library quality was finally assessed using Bioanalyzer high sensitivity DNA kit (Agilent). Paired-end sequencing was performed on a NextSeq 550 platform (Illumina).

Reads were quality checked using FastQC (available at: http://www.bioinformatics.bbsrc.ac.uk/projects/fastqc) and aligned to mm10 using Bowtie2 v2.4.2 (Langmead and Salzberg, 2012) with the following options: -X 2000 –dovetail. Unique reads (>q30) aligned to chromosomes 1–19 and chromosome X were retained. Peaks were called using MACS2 v2.2.7.1 (Zhang et al., 2008) with the following parameters: -q 0.01 – nomodel --shift −75 --extsize 150 -B –SPMR. Peaks called from individual samples were merged using mergePeaks.pl from the HOMER package v4.11 (Heinz et al., 2010) to generate a peak set on which to perform differential accessibility analyses. bedGraph files were converted into BigWig files using bedGraphtoBigWig and visualised in the UCSC Genome Browser (Kent et al., 2002). featureCounts v2.0.1 (Liao et al., 2014) was used to count reads within peaks from ATAC-seq samples and these were used an input for DESeq2 v1.34.0 to calculate differential binding using default settings. Counts were normalised using reads in peaks (RIP). A linear fold change of ±2 and a *q*-value of <0.05 were used as a cut-off for further analyses.

### ChIP-seq

Samples were prepared using iDeal ChIP-seq kit for Histones (Diagenode, C01010051). Briefly, 1×10^6^ *Fh1* ^*fl/fl*^ or 2×10^6^ *Fh1* ^*-/-*^ cells were plated onto 15-cm dishes (Nunc) and allowed to reach 80% confluency. Cells were then washed twice in PBS and detached using trypsin (Gibco, 0.25% in PBS-EDTA). After inactivating trypsin with complete medium, cells were filtered through 70μm cell strainers (Fisher Scientific) and counted with Cell Counter CASY (OmniLife Sciences). Following an additional wash in PBS, 7×10^6^ cells were resuspended in 500 μl of PBS and crosslinked using formaldehyde (Thermo Fisher) at 1% final concentration. Crosslinking was stopped using glycine solution after 10 min and 3 washes were carried in cold PBS before storing the pellets at -80°C. Further processing of the samples was carried out accordingly to the iDeal ChIP-seq kit for Histones protocol. 10-15 cycles of sonication (30 sec ON, 30 sec OFF) were carried using the Bioruptor Pico sonicator device (Diagenode). H3k27ac antibody was obtained from Abcam (ab4729) and used at the ratio 1μg/sheared chromatin derived from 1×10^6^ cells. Control rabbit IgG were purchased from Sigma. DNA Library was generated using NEB Next Ultra II DNA library kit (NEB, E7645S) following the manufacturer’s protocol. Library quality was assessed using Bioanalyzer high sensitivity DNA kit (Agilent). Single-end sequencing was performed on a NextSeq 550 platform (Illumina).

Reads were trimmed using Trimmomatic v0.34 (Bolger et al., 2014), quality checked using FastQC (available at: http://www.bioinformatics.bbsrc.ac.uk/projects/fastqc) and reads were aligned to mm10 and dm6 using Bowtie2 v2.4.2 (Langmead and Salzberg, 2012). Only reads with a mapping quality >q30 were retained. Peak calling was performed on merged replicates using MACS2 v2.2.7.1 (Zhang et al., 2008) using default parameters with additional –SPMR parameter and using the IgG sample as a control. bedGraph files were converted to bigwig using BedGraphtoBigWig script and visualised in the UCSC Genome Browser. featureCounts v2.0.1 (Liao et al., 2014) was used to count reads within peaks from ChIP-seq samples and these were used an input for DESeq2 v1.34.0 to calculate differential binding using default settings. Counts were normalised using reads in peaks (RIP). A linear fold change of ±2 and a *q*-value of <0.05 were used as a cut-off for further analyses.

### Data integration

The ATAC-seq peakset, originating from all cell-lines, was used as a reference. Peaks were separated into whether they showed a significant increase, decrease or no change between *Fh1*^*fl/fl*^ and *Fh1*^*-/-CL1*^ cells. Peaks were further separated into whether they overlapped a region displaying a significant increase, decrease or no change in H3K27ac ChIP-seq signal. Overlaps were calculated using intersect from BEDTools v2.30.0 (Quinlan and Hall, 2010) using parameters -wa -u to obtain positive overlaps, and parameters -v to obtain regions with no overlaps.

### Transcription factor motif analysis

To analyse multiple regions for enriched transcription factor motifs, genomic coordinates were analysed using gimme maelstrom from the GimmeMotifs v0.16.0 package (Bruse and Heeringen, 2018; van Heeringen and Veenstra, 2011) using the default motif database. To analyse single datasets, findMotifsGenome.pl from the HOMER package was used (Heinz et al., 2010) with –cpg –mask parameters.

### Footprinting analysis

To analyse footprinting signatures in ATAC-seq data the TOBIAS v0.12.12 package was used (Bentsen et al., 2020). Merged BAM files from each condition were processed using ATACorrect, footprint scores calculated using FootprintScores and differential footprinting analysis was performed using BINDetect using only regions identified from each cluster. The difference in footprinting between *Fh1*^*fl/fl*^ and *Fh1*^*-/-CL1*^ from each cluster was visualised in the heatmap.

### CRISPR screen analysis

Comparative pooled CRISPR screen data between *Fh1*^*fl/fl*^ and *Fh1*^*-/-CL1*^ was obtained from Valcarcel-Jiminez (2022). Volcano plots were plotted using MAGeCKFlute v1.14.0 (Wang et al., 2019).

### Patient survival analysis

Patient survival was calculated using the kidney renal papillary cell carcinoma mRNA TCGA dataset (KIRP) from kmplot.com (Nagy et al., 2021) using default settings.

### siRNA transfection

5 × 10^4^ *Fh1*^*fl/fl*^ and 1 × 10^5^ *Fh1*^*-/-CL1*^ and *Fh1*^*-/-CL19*^ cells were reverse transfected in 6-well plates with 50 pmol of either control siNT (Horizon, D-001810-10-05) or siFoxa2 (Horizon, L-043601-00-0005) siRNA using 2.5 μl RNAiMAX per well. Cells were incubated for 72 hours before either RNA or protein were extracted.

### Protein extraction and immunoblotting

Cells were lysed directly in cold RIPA buffer, supplemented with protease inhibitor cocktail and benzonase and were incubated on ice for at least 10 minutes. Lysate was transferred to a microcentrifuge tube and an aliquot was used in a BCA assay to determine protein concentration. LDS loading buffer was added to the lysate at a final 1x concentration and samples were heated to 70 °C for 10 minutes. Equal amounts of protein were loaded onto a 4-12% PAGE gel and then transferred to a nitrocellulose membrane. Total protein was stained for using and scanned. Membranes were blocked in a 5% BSA/TBS solution for 1 hour and incubated with anti-Foxa2 or anti-FH antibody overnight at 4 °C. IRDye secondary antibodies were incubated at room temperature for 1 hour and membranes were imaged on a Licor scanner. Membranes were washed in 1x TBS with 0.1% Tween-20.

### Cell growth assay

5 × 10^4^ cells were reverse transfected with 12.5 pmol siRNA (siNT or siFoxa2) and were allowed to settle at room temperature for up to an hour. Plates were transferred into an Incucyte® machine and confluence was monitored every six hours for 5 days. 9 images were taken per well per timepoint. Images were processed by Incucyte® Base Analysis Software using default settings.

### RNA-seq and analysis

RNA was extracted from cells using a RNeasy Plus RNA extraction Kit (Qiagen, 74136) and quality checked using Nanodrop 1000 (ThermoFisher). Single-end RNA-seq libraries were generated using TruSeq stranded mRNA library kit (Illumina) and sequenced on a NextSeq 550 platform (Illumina) by the Cambridge Genomics Services.

Reads were trimmed using Trimmomatic v0.39 (Bolger et al., 2014), quality checked using FastQC (available at: http://www.bioinformatics.bbsrc.ac.uk/projects/fastqc) and aligned to mm10 using STAR v2.7.9a using default parameters (Dobin et al., 2013). Reads aligned to chromosomes 1–19 and chromosome X were retained. Counts for genes were determined using featureCounts v2.0.1 (Liao et al., 2014) and DESeq2 v1.34.0 (Love et al., 2014) was used to perform differential gene expression analysis. Significant gene expression changes were defined by a fold change of ±2^0.5^ and a *q*-value of <0.05.

RNA-seq data from *Fh1*^*fl/fl*^, *Fh1*^*-/-CL1*^*+Fh1-GFP, Fh1*^*-/-CL1*^ and *Fh1*^*-/-CL19*^ cells were obtained from Sciacovelli et al., 2016 (GSE77542). RNA-seq data for FH-deficient tumours were obtained from Crooks et al., 2021 (GSE157256). To stratify TCGA KIRP (The Cancer Genome Atlas Research Network, 2016) tumour samples into FH positive and negative, samples were ranked according to FH expression, then the top 25% were classed as FH high and the bottom 25% were classed as FH low and then compared with DESeq2 v1.34.0 (Love et al., 2014).

### ChIPmentation and analysis

ChIPmentation was carried out as described previously (Schmidl et al., 2015). 1 × 10^6^ *Fh1*^*fl/fl*^ and 2 × 10^6^ *Fh1*^*-/-CL1*^ cells were cultured for 48 hours in 15 cm dish and crosslinked for 10 minutes in 1% formaldehyde. Cross-linking was quenches by 0.125 M glycine for at least 5 minutes. Cells were washed twice in 1 x PBS and scraped into ice-cold 1 x PBS with protease inhibitor cocktail and flash frozen and stored at -80°C until needed. 2.5 μg anti-FOXA2 antibody (abcam, ab256493) and 2.5 μg normal IgG antibody was used (Merck, 12-370). 1 μg Spike-In antibody (Active Motif, 61686) was also added to each replicate. 20 ng Spike-in Drosophila chromatin (Active Motif, 53083) was supplemented to chromatin preps for Spike-in normalisation, as described previously (Egan et al., 2016). DNA libraries were prepared using TruSeq ChIP sample prep kit (Illumina) and sequenced on a NextSeq 550 (Illumina) platform.

Reads were trimmed using Trimmomatic v0.34 (Bolger et al., 2014), quality checked using FastQC (available at: http://www.bioinformatics.bbsrc.ac.uk/projects/fastqc) and reads were aligned to mm10 and dm6 using Bowtie2 v2.4.2 (Langmead and Salzberg, 2012). Reads aligning to the Drosophila genome were counted and used to generate scale factors. BAM files were then scaled to the sample with the lowest number of Drosophila reads. Only reads with a mapping quality >q30 were retained. Peak calling was performed on merged replicates using MACS2 v2.2.7.1 (Zhang et al., 2008) using default parameters with additional –SPMR parameter and using the IgG sample as a control. bedGraph files were converted to bigwig using BedGraphtoBigWig script and visualised in the UCSC Genome Browser.

### Rapid immunoprecipitation mass spectrometry of endogenous proteins (RIME)

RIME was carried out as described previously reported (Glont et al., 2019; Papachristou et al., 2018). 2 × 10^6^ *Fh1*^*fl/fl*^ and 4 × 10^6^ *Fh1*^*-/-CL1*^ cells were cultured for 48 hours in 2 × 15 cm plates and were then dual crosslinked with 2mM DSG for 20 minutes, and then 1% formaldehyde for 10 minutes. Cross-linking was quenched by 0.125 M glycine for at least 5 minutes. Cells were washed twice in 1 x PBS and scraped into ice-cold 1 x PBS with protease inhibitor cocktail and flash frozen and stored at -80°C until needed. 5 μg anti-FOXA2 antibody (abcam, ab256493) and 5 μg normal IgG antibody was used.

Beads were digested with a concentration of 15 ng/μl Trypsin (Pierce) overnight at 37°C. The next day Trypsin was added again for a second 4 hr digestion at 37°C. The peptides were acidified with 5% formic acid, purified using the Ultra-Micro C18 Spin Columns (Harvard Apparatus), and dried with a vacuum concentrator. The samples were reconstituted in 0.1 % formic acid then analysed on a Dionex Ultimate 3000 UHPLC system coupled with a Q Exactive HF (Thermo Scientific) mass spectrometer. The full MS scans ran in the orbitrap with the range 400 – 1600 m/z at 60 K resolution. The 10 most intense precursors were selected for MS2 at resolution 30 K with an isolation window of 2.0 m/z and HCD collision energy at 28 %. The HCD spectra were processed with Proteome Discoverer 1.4 using the SequestHT search engine and a mouse uniprot database with over 17000 entries. The parameters for the analysis included: maximum of 2 missed cleavage sites, Precursor Mass Tolerance 20ppm, Fragment Mass Tolerance 0.02Da, and dynamic modifications deamidation of N/Q (+0.984Da) and Oxidation of M (+15.995Da). The percolator node was applied using a decoy database search. Peptides were filtered for a Target FDR with q-value<0.01. Specific interactors were from all three sample replicates but didn’t occur in the corresponding IgG injections. One of the *Fh1*^*-/-CL1*^ IgG samples was removed from the analysis as it was a clear outlier.

### Metabolomics and analysis

5 × 10^4^ *Fh1*^*fl/fl*^ and 1 × 10^5^ *Fh1*^*-/-CL1*^ and *Fh1*^*-/-CL19*^ cells were plated the onto 6-well plate and reverse transfected with siRNA. Before extraction, cells were counted using a separate counting plate. After that, cells were washed at room temperature with PBS twice and then kept on cold bath with dry ice and methanol. Metabolite extraction buffer (50% methanol, 30% acetonitrile, 20% ultrapure water, 5 μM final concentration valine-d8) was added to each well following the proportion 1×10^6^ cells/0.5 ml of buffer. Plates were kept at −80°C and kept overnight. The following day, the extracts were scraped and mixed at 4°C for 15 min. After final centrifugation at max speed for 10 min at 4°C, the supernatants were transferred into LC-MS vials.

Chromatographic separation of metabolites was achieved using a Millipore Sequant ZIC-pHILIC analytical column (5 μm, 2.1 × 150 mm) equipped with a 2.1 × 20 mm guard column (both 5 mm particle size) with a binary solvent system. Solvent A was 20 mM ammonium carbonate, 0.05% ammonium hydroxide; Solvent B was acetonitrile. The column oven and autosampler tray were held at 40 °C and 4 °C, respectively. The chromatographic gradient was run at a flow rate of 0.200 mL/min as follows: 0–2 min: 80% B; 2-17 min: linear gradient from 80% B to 20% B; 17-17.1 min: linear gradient from 20% B to 80% B; 17.1-23 min: hold at 80% B. Samples were randomized and the injection volume was 5 μl. A pooled quality control (QC) sample was generated from an equal mixture of all individual samples and analysed interspersed at regular intervals.

Metabolites were measured with Vanquish Horizon UHPLC coupled to an Orbitrap Exploris 240 mass spectrometer (both Thermo Fisher Scientific) via a heated electrospray ionization source. The spray voltages were set to +3.5kV/-2.8 kV, RF lens value at 70, the heated capillary held at 320 °C, and the auxiliary gas heater held at 280 °C. The flow rate for sheath gas, aux gas and sweep gas were set to 40, 15 and 0, respectively. For MS1 scans, mass range was set to m/z=70-900, AGC target set to standard and maximum injection time (IT) set to auto. Data acquisition for experimental samples used full scan mode with polarity switching at an Orbitrap resolution of 120000. Data acquisition for untargeted metabolite identification was performed using the AcquireX Deep Scan workflow, an iterative data-dependent acquisition (DDA) strategy using multiple injections of the pooled sample. In brief, sample was first injected in full scan-only mode in single polarity to create an automated inclusion list. MS2 acquisition was then carried out in triplicate, where ions on the inclusion list were prioritized for fragmentation in each run, after which both the exclusion and inclusion lists were updated in a manner where fragmented ions from the inclusion list were moved to exclusion list for the next run. DDA full scan-ddMS2 method for AcquireX workflow used the following parameters: full scan resolution was set to 60000, fragmentation resolution to 30000, fragmentation intensity threshold to 5.0e3. Dynamic exclusion was enabled after 1 time and exclusion duration was 10s. Mass tolerance was set to 5ppm. Isolation window was set to 1.2 m/z. Normalized HCD collision energies were set to stepped mode with values at 30, 50, 150. Fragmentation scan range was set to auto, AGC target at standard and max IT at auto. Xcalibur AcquireX method modification was on. Mild trapping was enabled.

Metabolite identification was performed in the Compound Discoverer software (v 3.2, Thermo Fisher Scientific). Metabolites were annotated at the MS2 level using both an in-house mzVault spectral database curated from 1051 authentic compound standards and the online spectral library mzCloud. The precursor mass tolerance was set to 5 ppm and fragment mass tolerance set to 10 ppm. Only metabolites with mzVault or mzCloud best match score above 50% and 75%, respectively, and RT tolerance within 0.5 min to that of a purified standard run with the same chromatographic method were exported to generate a list including compound names, molecular formula and RT. The curated list was then used for further processing in the Tracefinder software (v 5.0, Thermo Fisher Scientific), where extracted ion chromatographs for all compound were examined and manually integrated if necessary. False positive, noise or chromatographically unresolved compounds were removed. The peak area for each detected metabolite was then normalized against the total ion count (TIC) of that sample to correct any variations introduced from sample handling through instrument analysis. The normalized areas were used as variables for further statistical data analysis.

### Data analysis and visualisation

All visualisation of ATAC-seq and ChIP-seq was done using the deepTools2 v3.5.1 package (Ramírez et al., 2016). ATAC-seq fragment size was visualised using bamPEFragmentSize. Correlation plots between technical replicates were visualised using multiBamSummary and plotCorrelation. Tag density plots and heatmaps were generated using computeMatrix and plotProfile or plotHeatmap tools.

Euler diagrams were generated using eulerr.co. Heatmaps were generated using the ComplexHeatmap v2.10.0 R package (Gu et al., 2016) and volcano plots were generated using the enhancedVolcano v1.12.0 R package (Blighe et al., 2018).

Peaks were assigned to their closest TSS/gene using the seq2gene function from the ChIPSeeker v1.30.3 R package (Yu et al., 2015). GO term enrichment was performed using the clusterProfiler v4.2.2 R package (Wu et al., 2021). Gene Set Enrichment Analysis was performed using the fgsea v1.20.0 R package (Korotkevich et al., 2021). Associating genomic regions to genes was done using PEGS v0.6.4 (Briggs et al., 2021). Random gene sets were generated using https://molbiotools.com/randomgenesetgenerator.php.

*P*-values were calculated in GraphPad Prism v9.0 using the student’s T-test with Welch’s correction, unless otherwise stated.

### Code

Code and scripts to reproduce the analyses are available at https://github.com/connorrogerson/FH_Foxa2 and https://github.com/ArianeMora/foxa2_kirp_kirc

### Data availability

All sequencing data is available from ArrayExpress (https://www.ebi.ac.uk/arrayexpress/) using the following identifiers: E-MTAB-11789 for ATAC-seq data; E-MTAB-11790 for the H3K27ac ChIP-seq data; E-MTAB-11782 for the siFoxa2 RNA-seq data and E-MTAB-11785 for the FOXA2 ChIPmentation data. The proteomics data is available at the PRIDE repository (https://www.ebi.ac.uk/pride/) with the dataset identifier PXD034141. The siFoxa2 metabolomics is available at Metabolomics Workbench (https://www.metabolomicsworkbench.org) under the accession ST002199.

## Acknowledgements

We thank Cambridge Genomic Services (Department of Pathology, University of Cambridge) especially Dr Alexandria Karcanias and Dr Julien Bauer for the RNA-seq library preparation and sequencing, and the Babraham Sequencing Facility for sequencing ChIP-seq and ATAC-seq libraries. We also thank all the Frezza lab members for their helpful discussions and insights, and the MRC Cancer Unit facilities and media kitchen for their invaluable support.

C.R. was supported by the ERC Consolidator Grant (ONCOFUM, ERC819920) to C.F. C.F is supported by the Medical Research Council (MRC_MC_UU_12022/6), the CRUK Programme Foundation award (C51061/A27453), ERC Consolidator Grant (ONCOFUM, ERC819920) and by the Alexander von Humboldt Foundation in the framework of the Alexander von Humboldt Professorship endowed by the Federal Ministry of Education and Research.

L V-J is supported by the long-term FEBS fellowship and by the CRUK Programme Foundation award to C.F. (C51061/A27453). C.S. was funded by the European Union’s Horizon 2020 research and innovation programme under the Marie Skłodowska-Curie grant agreement #722605 and the Alexander von Humboldt Professorship to C.F. A.M. was funded by Australian Government Research Training Program.

G.K. is supported by the UK Biotechnology and Biological Sciences Research Council (BBS/E/B/000C0423) and Medical Research Council (MR/S000437/1). J.S.C. is supported by Cancer Research UK core funding (A20411 and A31344).

## Author contributions

Conceptualisation, C.R. and C.F.; Formal Analysis, C.R., C.S., M.Y., J.K. and A.M.; Investigation, C.R., M.S., L.A.M., L.V-J., E.I. and J.K.; Resources, D.C and J.S.C.; Writing – Original Draft, C.R.; Writing – Review & Editing, C.R. M.S., L.A.M., L.V-J., C.S., J.S.C, G.K. and C.F.; Visualisation, C.R.; Supervision, J.S.C., G.K., C.F.; Project administration, C.F.; Funding Acquisition, C.F.

## Figure Legends

**Supplementary Figure 1**

(A) Histogram of fragment length of ATAC-seq sequencing data from *Fh1*^*fl/fl*^, *Fh1*^*-/-CL1*^*+pFh1-GFP, Fh1*^*-/-CL1*^, *Fh1*^*-/-CL19*^ replicates. (B) Pearson correlation of ATAC-seq replicate data from *Fh1*^*fl/fl*^, *Fh1*^*-/-CL1*^*+pFh1-GFP, Fh1*^*-/-CL1*^, *Fh1*^*-/-CL19*^ cells (C) Pearson correlation of ChIP-seq replicate data from *Fh1*^*fl/fl*^, *Fh1*^*-/-CL1*^*+pFh1-GFP, Fh1*^*-/-CL1*^, *Fh1*^*-/-CL19*^ cells. (D) Normalised H3K27ac ChIP-seq and ATAC-seq signal at the *Mir200c* (upper) and Slc7a11 (lower) loci. (E) Normalised ATAC-seq (upper) and H3K27ac ChIP-seq signal (lower) from from *Fh1*^*fl/fl*^ and *Fh1*^*-/-CL19*^ cells at differentially accessible and differentially bound regions.

**Supplementary Figure 2**

(A) Heatmap of normalised ATAC-seq signal at clusters I – VII. (B) Heatmap of normalised H3K27ac ChIP-seq signal at clusters I – VII. (C) Bar chart of the percentage of regions within intervals of distance from the nearest transcription start site (TSS). (D) Heatmap of enrichment *P*-values (hypergeometric test) of differentially expressed genes in *Fh1*^*-/-CL1*^ compared to *Fh1*^*fl/fl*^ associated with clusters I - VII at increasing distances from the TSS.

**Supplementary Figure 3**

(A) Expression of forkhead factors in *Fh1*^*-/-CL1*^ cells ranked from lowest expression to the highest. (B) Log_2_(Fold change) of significantly differentially expressed forkhead factors in *Fh1*^*-/-CL1*^ compared to *Fh1*^*fl/fl*^ cells. (C) Box plot of *FOXA2* expression in human FH-deficient papillary renal cell carcinoma compared to adjacent normal tissue (Crooks et al., 2021). (D) Box plot of *FOXA2* expression in FH-high and FH-low papillary renal cell carcinoma tissue (TCGA KIRP) (E) Kaplan-Meier curves of overall patient survival for high (above median; red) or low (below median; black) expression of *FOXA2* in renal papillary cell carcinoma.

**Supplementary Figure 4**

(A) Immunoblot of protein lysate from *Fh1*^*fl/fl*^, *Fh1*^*-/-CL1*^ and *Fh1*^*-/-CL19*^ cells treated with either siNT or siFoxa2 probed with antibodies against FOXA2. (B) Pearson correlation of RNA-seq replicate data from *Fh1*^*fl/fl*^, *Fh1*^*-/-CL1*^ and *Fh1*^*-/-CL19*^ cells treated with either siNT or siFoxa2. (C) Euler diagram of DEGs from *Fh1*^*fl/fl*^, *Fh1*^*-/-CL1*^ and *Fh1*^*-/-CL19*^ cells treated with either siFoxa2 compared to siNT. (D) Heatmap of enrichment *P*-values (hypergeometric test) of differentially expressed genes from siFoxa2 treatment associated with clusters I - VII at increasing distances from the TSS.

**Supplementary Figure 5**

(A) Scatter plot of FOXA2 ChIP-seq signal between replicates in *Fh1*^*fl/fl*^ cells. R^2^ value shown. (B) Scatter plot of FOXA2 ChIP-seq signal between replicates in *Fh1*^*-/-CL1*^ cells. R^2^ value shown. (C) Normalised ATAC-seq signal (upper) and H3K27ac ChIP-seq signal (lower) from *Fh1*^*fl/fl*^ (blue), *Fh1*^*-/-CL1*^ (red) cells at *Fh1*^*-/-CL1*^ specific peaks (left), shared peaks (middle) and *Fh1*^*fl/fl*^ specific peaks (right). (D) Heatmap of enrichment *P*-values (hypergeometric test) of NRF2 target genes (NFE2L2.V2 gene set) to FOXA2 ChIP-seq peaks at increasing distances from the TSS. (E) Line plot of normalised abundances of FOXA2, AP-1, NRF2 and CTCF motifs across *Fh1*^*-/-CL1*^ specific peaks.

**Supplementary Figure 6**

PCA plot of normalised metabolomics replicates from *Fh1*^*fl/fl*^, *Fh1*^*-/-CL1*^, *Fh1*^*-/-CL19*^ cells treated with either siNT or siFoxa2.

## References

Adam, J., Hatipoglu, E., O’Flaherty, L., Ternette, N., Sahgal, N., Lockstone, H., Baban, D., Nye, E., Stamp, G.W., Wolhuter, K., Stevens, M., Fischer, R., Carmeliet, P., Maxwell, P.H., Pugh, C.W., Frizzell, N., Soga, T., Kessler, B.M., El-Bahrawy, M., Ratcliffe, P.J., Pollard, P.J., 2011. Renal Cyst Formation in Fh1-Deficient Mice Is Independent of the Hif/Phd Pathway: Roles for Fumarate in KEAP1 Succination and Nrf2 Signaling. Cancer Cell 20, 524–537. https://doi.org/10.1016/j.ccr.2011.09.006

Alderson, N.L., Wang, Y., Blatnik, M., Frizzell, N., Walla, M.D., Lyons, T.J., Alt, N., Carson, J.A., Nagai, R., Thorpe, S.R., Baynes, J.W., 2006. S-(2-Succinyl)cysteine: A novel chemical modification of tissue proteins by a Krebs cycle intermediate. Archives of Biochemistry and Biophysics 450, 1–8. https://doi.org/10.1016/j.abb.2006.03.005

Baird, L., Yamamoto, M., 2020. The Molecular Mechanisms Regulating the KEAP1-NRF2 Pathway. Molecular and Cellular Biology 40, e00099–20. https://doi.org/10.1128/MCB.00099-20

Bentsen, M., Goymann, P., Schultheis, H., Klee, K., Petrova, A., Wiegandt, R., Fust, A., Preussner, J., Kuenne, C., Braun, T., Kim, J., Looso, M., 2020. ATAC-seq footprinting unravels kinetics of transcription factor binding during zygotic genome activation. Nat Commun 11, 4267. https://doi.org/10.1038/s41467-020-18035-1

Blighe, K., Rana, S., Lewis, M, 2018. EnhancedVolcano: publication-ready volcano plots with enhanced colouring and labeling.

Boivin, F.J., Schmidt-Ott, K.M., 2020. Functional roles of Grainyhead-like transcription factors in renal development and disease. Pediatr Nephrol 35, 181–190. https://doi.org/10.1007/s00467-018-4171-4

Bolger, A.M., Lohse, M., Usadel, B., 2014. Trimmomatic: a flexible trimmer for Illumina sequence data. Bioinformatics 30, 2114–2120. https://doi.org/10.1093/bioinformatics/btu170

Briggs, P., Hunter, A., Yang, S., Sharrocks, A., Iqbal, M., 2021. PEGS: An efficient tool for gene set enrichment within defined sets of genomic intervals. F1000Research 10. https://doi.org/10.12688/f1000research.53926.2

Bruse, N., Heeringen, S.J. van, 2018. GimmeMotifs: an analysis framework for transcription factor motif analysis. bioRxiv 474403. https://doi.org/10.1101/474403

Corces, M.R., Trevino, A.E., Hamilton, E.G., Greenside, P.G., Sinnott-Armstrong, N.A., Vesuna, S., Satpathy, A.T., Rubin, A.J., Montine, K.S., Wu, B., Kathiria, A., Cho, S.W., Mumbach, M.R., Carter, A.C., Kasowski, M., Orloff, L.A., Risca, V.I., Kundaje, A., Khavari, P.A., Montine, T.J., Greenleaf, W.J., Chang, H.Y., 2017. An improved ATAC-seq protocol reduces background and enables interrogation of frozen tissues. Nature Methods 14, 959–962. https://doi.org/10.1038/nmeth.4396

Creyghton, M.P., Cheng, A.W., Welstead, G.G., Kooistra, T., Carey, B.W., Steine, E.J., Hanna, J., Lodato, M.A., Frampton, G.M., Sharp, P.A., Boyer, L.A., Young, R.A., Jaenisch, R., 2010. Histone H3K27ac separates active from poised enhancers and predicts developmental state. Proceedings of the National Academy of Sciences 107, 21931–21936. https://doi.org/10.1073/pnas.1016071107

Crooks, D.R., Maio, N., Lang, M., Ricketts, C.J., Vocke, C.D., Gurram, S., Turan, S., Kim, Y.-Y., Cawthon, G.M., Sohelian, F., De Val, N., Pfeiffer, R.M., Jailwala, P., Tandon, M., Tran, B., Fan, T.W.-M., Lane, A.N., Ried, T., Wangsa, D., Malayeri, A.A., Merino, M.J., Yang, Y., Meier, J.L., Ball, M.W., Rouault, T.A., Srinivasan, R., Linehan, W.M., 2021. Mitochondrial DNA alterations underlie an irreversible shift to aerobic glycolysis in fumarate hydratase–deficient renal cancer. Science Signaling 14, eabc4436. https://doi.org/10.1126/scisignal.abc4436

DeBlasi, J.M., DeNicola, G.M., 2020. Dissecting the Crosstalk between NRF2 Signaling and Metabolic Processes in Cancer. Cancers 12, 3023. https://doi.org/10.3390/cancers12103023

Dobin, A., Davis, C.A., Schlesinger, F., Drenkow, J., Zaleski, C., Jha, S., Batut, P., Chaisson, M., Gingeras, T.R., 2013. STAR: ultrafast universal RNA-seq aligner. Bioinformatics 29, 15–21. https://doi.org/10.1093/bioinformatics/bts635

Egan, B., Yuan, C.-C., Craske, M.L., Labhart, P., Guler, G.D., Arnott, D., Maile, T.M., Busby, J., Henry, C., Kelly, T.K., Tindell, C.A., Jhunjhunwala, S., Zhao, F., Hatton, C., Bryant, B.M., Classon, M., Trojer, P., 2016. An Alternative Approach to ChIP-Seq Normalization Enables Detection of Genome-Wide Changes in Histone H3 Lysine 27 Trimethylation upon EZH2 Inhibition. PLoS One 11. https://doi.org/10.1371/journal.pone.0166438

Frezza, C., Zheng, L., Folger, O., Rajagopalan, K.N., MacKenzie, E.D., Jerby, L., Micaroni, M., Chaneton, B., Adam, J., Hedley, A., Kalna, G., Tomlinson, I.P.M., Pollard, P.J., Watson, D.G., Deberardinis, R.J., Shlomi, T., Ruppin, E., Gottlieb, E., 2011. Haem oxygenase is synthetically lethal with the tumour suppressor fumarate hydratase. Nature 477, 225–228. https://doi.org/10.1038/nature10363

Friedmann-Morvinski, D., Verma, I.M., 2014. Dedifferentiation and reprogramming: origins of cancer stem cells. EMBO Rep 15, 244–253. https://doi.org/10.1002/embr.201338254

Gao, N., Le Lay, J., Qin, W., Doliba, N., Schug, J., Fox, A.J., Smirnova, O., Matschinsky, F.M., Kaestner, K.H., 2010. Foxa1 and Foxa2 Maintain the Metabolic and Secretory Features of the Mature β-Cell. Molecular Endocrinology 24, 1594–1604. https://doi.org/10.1210/me.2009-0513

Glont, S.-E., Chernukhin, I., Carroll, J.S., 2019. Comprehensive Genomic Analysis Reveals that the Pioneering Function of FOXA1 Is Independent of Hormonal Signaling. Cell Rep 26, 2558–2565.e3. https://doi.org/10.1016/j.celrep.2019.02.036

Grubb, R.L., Franks, M.E., Toro, J., Middelton, L., Choyke, L., Fowler, S., Torres, -Cabala Carlos, Glenn, G.M., Choyke, P., Merino, M.J., Zbar, B., Pinto, P.A., Srinivasan, R., Coleman, J.A., Linehan, W.M., 2007. Hereditary Leiomyomatosis and Renal Cell Cancer: A Syndrome Associated With an Aggressive Form of Inherited Renal Cancer. Journal of Urology 177, 2074–2080. https://doi.org/10.1016/j.juro.2007.01.155

Gu, Z., Eils, R., Schlesner, M., 2016. Complex heatmaps reveal patterns and correlations in multidimensional genomic data. Bioinformatics 32, 2847–2849. https://doi.org/10.1093/bioinformatics/btw313

Heinz, S., Benner, C., Spann, N., Bertolino, E., Lin, Y.C., Laslo, P., Cheng, J.X., Murre, C., Singh, H., Glass, C.K., 2010. Simple combinations of lineage-determining transcription factors prime cis-regulatory elements required for macrophage and B cell identities. Mol. Cell 38, 576–589. https://doi.org/10.1016/j.molcel.2010.05.004

Iwafuchi-Doi, M., Donahue, G., Kakumanu, A., Watts, J.A., Mahony, S., Pugh, B.F., Lee, D., Kaestner, K.H., Zaret, K.S., 2016. The Pioneer Transcription Factor FoxA Maintains an Accessible Nucleosome Configuration at Enhancers for Tissue-Specific Gene Activation. Mol. Cell 62, 79–91. https://doi.org/10.1016/j.molcel.2016.03.001

Jiang, Y., Qian, X., Shen, J., Wang, Y., Li, X., Liu, R., Xia, Y., Chen, Q., Peng, G., Lin, S.-Y., Lu, Z., 2015. Local generation of fumarate promotes DNA repair through inhibition of histone H3 demethylation. Nat Cell Biol 17, 1158–1168. https://doi.org/10.1038/ncb3209

Kent, W.J., Sugnet, C.W., Furey, T.S., Roskin, K.M., Pringle, T.H., Zahler, A.M., Haussler, and D., 2002. The Human Genome Browser at UCSC. Genome Res. 12, 996–1006. https://doi.org/10.1101/gr.229102

Korotkevich, G., Sukhov, V., Budin, N., Shpak, B., Artyomov, M.N., Sergushichev, A., 2021. Fast gene set enrichment analysis. bioRxiv 060012. https://doi.org/10.1101/060012

Kulkarni, R.A., Bak, D.W., Wei, D., Bergholtz, S.E., Briney, C.A., Shrimp, J.H., Alpsoy, A., Thorpe, A.L., Bavari, A.E., Crooks, D.R., Levy, M., Florens, L., Washburn, M.P., Frizzell, N., Dykhuizen, E.C., Weerapana, E., Linehan, W.M., Meier, J.L., 2019. A chemoproteomic portrait of the oncometabolite fumarate. Nat Chem Biol 15, 391– 400. https://doi.org/10.1038/s41589-018-0217-y

Langmead, B., Salzberg, S.L., 2012. Fast gapped-read alignment with Bowtie 2. Nature Methods 9, 357.

Laukka, T., Mariani, C.J., Ihantola, T., Cao, J.Z., Hokkanen, J., Kaelin, W.G., Godley, L.A., Koivunen, P., 2016. Fumarate and Succinate Regulate Expression of Hypoxia-inducible Genes via TET Enzymes. Journal of Biological Chemistry 291, 4256–4265. https://doi.org/10.1074/jbc.M115.688762

Li, M., Chiu, J.-F., Kelsen, A., Lu, S.C., Fukagawa, N.K., 2009. Identification and characterization of an Nrf2-mediated ARE upstream of the rat glutamate cysteine ligase catalytic subunit gene (GCLC). Journal of Cellular Biochemistry 107, 944–954. https://doi.org/10.1002/jcb.22197

Liao, Y., Smyth, G.K., Shi, W., 2014. featureCounts: an efficient general purpose program for assigning sequence reads to genomic features. Bioinformatics 30, 923–930. https://doi.org/10.1093/bioinformatics/btt656

Love, M.I., Huber, W., Anders, S., 2014. Moderated estimation of fold change and dispersion for RNA-seq data with DESeq2. Genome Biol 15. https://doi.org/10.1186/s13059-014-0550-8

Malhotra, D., Portales-Casamar, E., Singh, A., Srivastava, S., Arenillas, D., Happel, C., Shyr, C., Wakabayashi, N., Kensler, T.W., Wasserman, W.W., Biswal, S., 2010. Global mapping of binding sites for Nrf2 identifies novel targets in cell survival response through ChIP-Seq profiling and network analysis. Nucleic Acids Research 38, 5718– 5734. https://doi.org/10.1093/nar/gkq212

Menko, F.H., Maher, E.R., Schmidt, L.S., Middelton, L.A., Aittomäki, K., Tomlinson, I., Richard, S., Linehan, W.M., 2014. Hereditary leiomyomatosis and renal cell cancer (HLRCC): renal cancer risk, surveillance and treatment. Familial Cancer 13, 637– 644. https://doi.org/10.1007/s10689-014-9735-2

Nagy, Á., Munkácsy, G., Győrffy, B., 2021. Pancancer survival analysis of cancer hallmark genes. Sci Rep 11, 6047. https://doi.org/10.1038/s41598-021-84787-5

Ooi, A., Wong, J.-C., Petillo, D., Roossien, D., Perrier-Trudova, V., Whitten, D., Min, B.W.H., Tan, M.-H., Zhang, Z., Yang, X.J., Zhou, M., Gardie, B., Molinié, V., Richard, S., Tan, P.H., Teh, B.T., Furge, K.A., 2011. An Antioxidant Response Phenotype Shared between Hereditary and Sporadic Type 2 Papillary Renal Cell Carcinoma. Cancer Cell 20, 511–523. https://doi.org/10.1016/j.ccr.2011.08.024

Papachristou, E.K., Kishore, K., Holding, A.N., Harvey, K., Roumeliotis, T.I., Chilamakuri, C.S.R., Omarjee, S., Chia, K.M., Swarbrick, A., Lim, E., Markowetz, F., Eldridge, M., Siersbaek, R., D’Santos, C.S., Carroll, J.S., 2018. A quantitative mass spectrometry-based approach to monitor the dynamics of endogenous chromatin-associated protein complexes. Nat Commun 9, 2311. https://doi.org/10.1038/s41467-018-04619-5

Quinlan, A.R., Hall, I.M., 2010. BEDTools: a flexible suite of utilities for comparing genomic features. Bioinformatics 26, 841–842. https://doi.org/10.1093/bioinformatics/btq033

Ramírez, F., Ryan, D.P., Grüning, B., Bhardwaj, V., Kilpert, F., Richter, A.S., Heyne, S., Dündar, F., Manke, T., 2016. deepTools2: a next generation web server for deep-sequencing data analysis. Nucleic Acids Res 44, W160–W165. https://doi.org/10.1093/nar/gkw257

Reddy, N.M., Kleeberger, S.R., Yamamoto, M., Kensler, T.W., Scollick, C., Biswal, S., Reddy, S.P., 2007. Genetic dissection of the Nrf2-dependent redox signaling-regulated transcriptional programs of cell proliferation and cytoprotection. Physiol Genomics 32, 74–81. https://doi.org/10.1152/physiolgenomics.00126.2007

Ryan, D.G., Yang, M., Prag, H.A., Blanco, G.R., Nikitopoulou, E., Segarra-Mondejar, M., Powell, C.A., Young, T., Burger, N., Miljkovic, J.L., Minczuk, M., Murphy, M.P., von Kriegsheim, A., Frezza, C., 2021. Disruption of the TCA cycle reveals an ATF4-dependent integration of redox and amino acid metabolism. eLife 10, e72593. https://doi.org/10.7554/eLife.72593

Schmidl, C., Rendeiro, A.F., Sheffield, N.C., Bock, C., 2015. ChIPmentation: fast, robust, low-input ChIP-seq for histones and transcription factors. Nat Methods 12, 963–965. https://doi.org/10.1038/nmeth.3542

Schmidt, C., Sciacovelli, M., Frezza, C., 2020. Fumarate hydratase in cancer: A multifaceted tumour suppressor. Seminars in Cell & Developmental Biology, SI: Cancer Cells & Therapeutic Targets 98, 15–25. https://doi.org/10.1016/j.semcdb.2019.05.002

Sciacovelli, M., Gonçalves, E., Johnson, T.I., Zecchini, V.R., da Costa, A.S.H., Gaude, E., Drubbel, A.V., Theobald, S.J., Abbo, S.R., Tran, M.G.B., Rajeeve, V., Cardaci, S., Foster, S., Yun, H., Cutillas, P., Warren, A., Gnanapragasam, V., Gottlieb, E., Franze, K., Huntly, B., Maher, E.R., Maxwell, P.H., Saez-Rodriguez, J., Frezza, C., 2016. Fumarate is an epigenetic modifier that elicits epithelial-to-mesenchymal transition. Nature 537, 544–547. https://doi.org/10.1038/nature19353

Sun, G., Zhang, X., Liang, J., Pan, X., Zhu, S., Liu, Z., Armstrong, C.M., Chen, Jianhui, Lin, W., Liao, B., Lin, T., Huang, R., Zhang, M., Zheng, L., Yin, X., Nie, L., Shen, P., Zhao, J., Zhang, H., Dai, J., Shen, Y., Li, Z., Liu, Jiyan, Chen, Junru, Liu, Jiandong, Wang, Z., Zhu, X., Ni, Y., Qin, D., Yang, L., Chen, Y., Wei, Q., Li, X., Zhou, Q., Huang, H., Yao, J., Chen, N., Zeng, H., 2021. Integrated Molecular Characterization of Fumarate Hydratase–deficient Renal Cell Carcinoma. Clinical Cancer Research 27, 1734–1743. https://doi.org/10.1158/1078-0432.CCR-20-3788

The Cancer Genome Atlas Research Network, 2016. Comprehensive Molecular Characterization of Papillary Renal-Cell Carcinoma. New England Journal of Medicine 374, 135–145. https://doi.org/10.1056/NEJMoa1505917

Thiery, J.P., Acloque, H., Huang, R.Y.J., Nieto, M.A., 2009. Epithelial-Mesenchymal Transitions in Development and Disease. Cell 139, 871–890. https://doi.org/10.1016/j.cell.2009.11.007

Tomlinson, I.P.M., Alam, N.A., Rowan, A.J., Barclay, E., Jaeger, E.E.M., Kelsell, D., Leigh, I., Gorman, P., Lamlum, H., Rahman, S., Roylance, R.R., Olpin, S., Bevan, S., Barker, K., Hearle, N., Houlston, R.S., Kiuru, M., Lehtonen, R., Karhu, A., Vilkki, S., Laiho, P., Eklund, C., Vierimaa, O., Aittomäki, K., Hietala, M., Sistonen, P., Paetau, A., Salovaara, R., Herva, R., Launonen, V., Aaltonen, L.A., The Multiple Leiomyoma Consortium, Group 1, Group 2, Group 3, 2002. Germline mutations in FH predispose to dominantly inherited uterine fibroids, skin leiomyomata and papillary renal cell cancer. Nature Genetics 30, 406–410. https://doi.org/10.1038/ng849

Valcarcel-Jimenez, L., Rogerson, C., Yong, C., Schmidt, C., Yang, M., Harle, V., Offord, V., Wong, K., Mora, A., Speed, A., Caraffini, V., Tran, M.G.B., Maher, E.R., Stewart, G.D., Vanharanta, S., Adams, D.J., Frezza, C., 2022. HIRA loss transforms FH-deficient cells. bioRxiv 2022.06.04.492123. https://doi.org/10.1101/2022.06.04.492123

van Heeringen, S.J., Veenstra, G.J.C., 2011. GimmeMotifs: a de novo motif prediction pipeline for ChIP-sequencing experiments. Bioinformatics 27, 270–271. https://doi.org/10.1093/bioinformatics/btq636

Wang, B., Wang, M., Zhang, W., Xiao, T., Chen, C.-H., Wu, A., Wu, F., Traugh, N., Wang, X., Li, Z., Mei, S., Cui, Y., Shi, S., Lipp, J.J., Hinterndorfer, M., Zuber, J., Brown, M., Li, W., Liu, X.S., 2019. Integrative analysis of pooled CRISPR genetic screens using MAGeCKFlute. Nat Protoc 14, 756–780. https://doi.org/10.1038/s41596-018-0113-7

Wolfrum, C., Asilmaz, E., Luca, E., Friedman, J.M., Stoffel, M., 2004. Foxa2 regulates lipid metabolism and ketogenesis in the liver during fasting and in diabetes. Nature 432, 1027–1032. https://doi.org/10.1038/nature03047

Xiao, M., Yang, H., Xu, W., Ma, S., Lin, H., Zhu, H., Liu, L., Liu, Y., Yang, C., Xu, Y., Zhao, S., Ye, D., Xiong, Y., Guan, K.-L., 2012. Inhibition of α-KG-dependent histone and DNA demethylases by fumarate and succinate that are accumulated in mutations of FH and SDH tumor suppressors. Genes Dev. 26, 1326–1338. https://doi.org/10.1101/gad.191056.112

Yu, G., Wang, L.-G., He, Q.-Y., 2015. ChIPseeker: an R/Bioconductor package for ChIP peak annotation, comparison and visualization. Bioinformatics 31, 2382–2383. https://doi.org/10.1093/bioinformatics/btv145

Zaret, K.S., Carroll, J.S., 2011. Pioneer transcription factors: establishing competence for gene expression. Genes Dev. 25, 2227–2241. https://doi.org/10.1101/gad.176826.111

Zhang, L., Rubins, N.E., Ahima, R.S., Greenbaum, L.E., Kaestner, K.H., 2005. Foxa2 integrates the transcriptional response of the hepatocyte to fasting. Cell Metabolism 2, 141–148. https://doi.org/10.1016/j.cmet.2005.07.002

Zhang, P., Sun, Y., Ma, L., 2015. ZEB1: At the crossroads of epithelial-mesenchymal transition, metastasis and therapy resistance. Cell Cycle 14, 481–487. https://doi.org/10.1080/15384101.2015.1006048

Zhang, Y., Liu, T., Meyer, C.A., Eeckhoute, J., Johnson, D.S., Bernstein, B.E., Nusbaum, C., Myers, R.M., Brown, M., Li, W., Liu, X.S., 2008. Model-based Analysis of ChIP-Seq (MACS). Genome Biology 9, R137. https://doi.org/10.1186/gb-2008-99-r137

Zheng, L., Cardaci, S., Jerby, L., MacKenzie, E.D., Sciacovelli, M., Johnson, T.I., Gaude, E., King, A., Leach, J.D.G., Edrada-Ebel, R., Hedley, A., Morrice, N.A., Kalna, G., Blyth, K., Ruppin, E., Frezza, C., Gottlieb, E., 2015. Fumarate induces redox-dependent senescence by modifying glutathione metabolism. Nat Commun 6, 6001. https://doi.org/10.1038/ncomms7001

