## Supplementary figures and images for "FOXA2 controls the anti-oxidant response in FH-deficient cells"

### Supplementary Figure 1

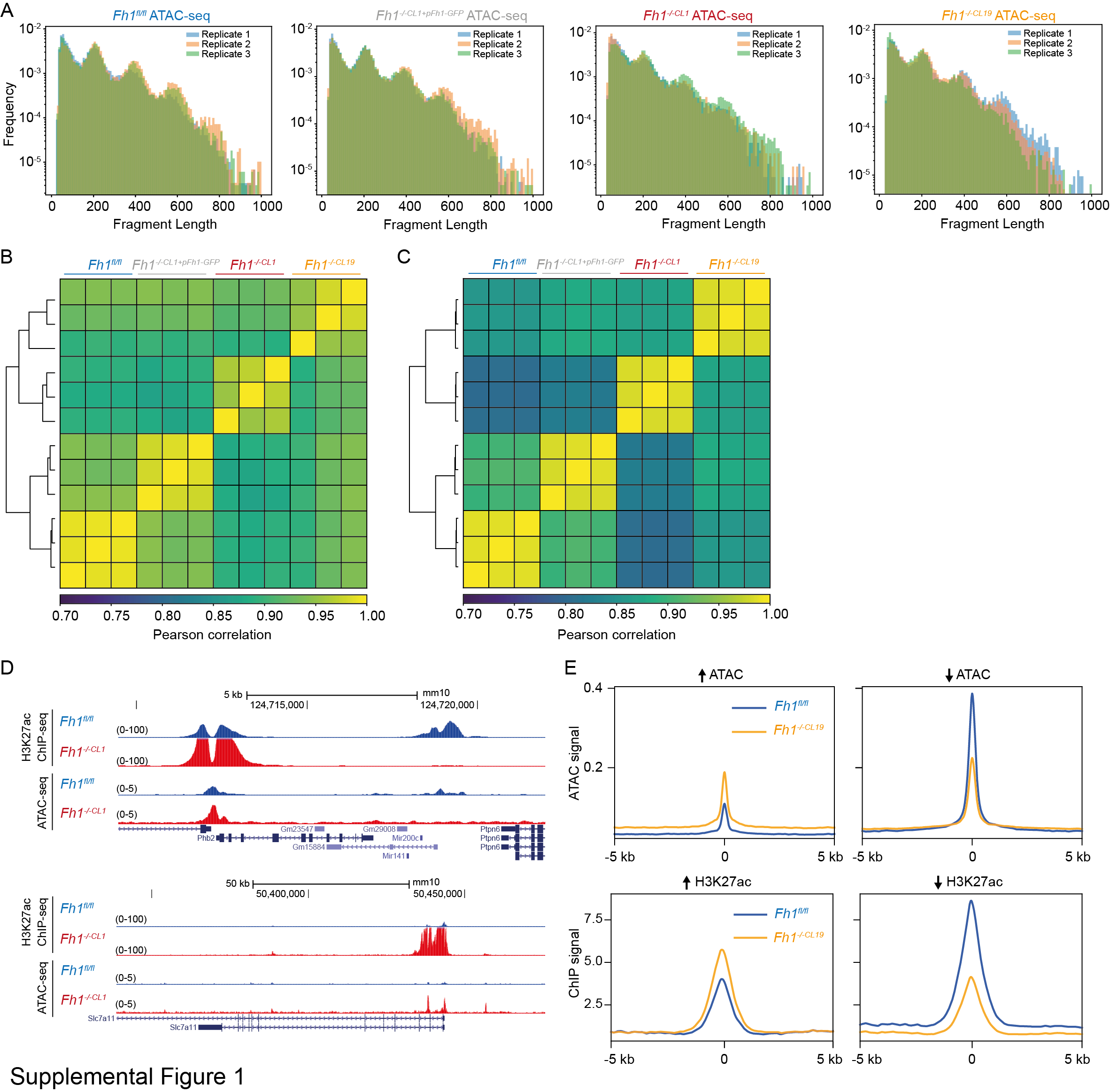

### Supplementary Figure 2

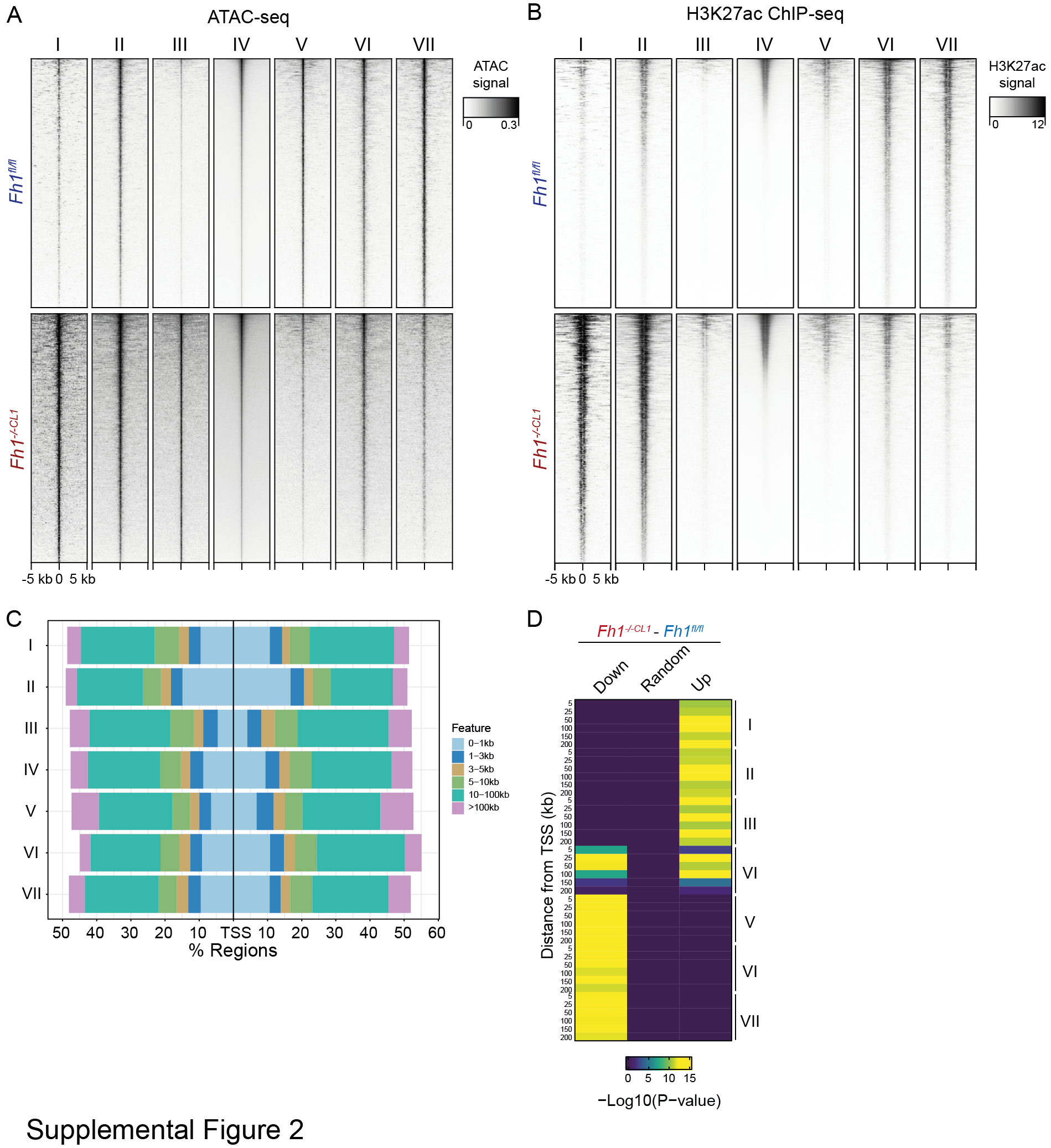

### Supplementary Figure 3

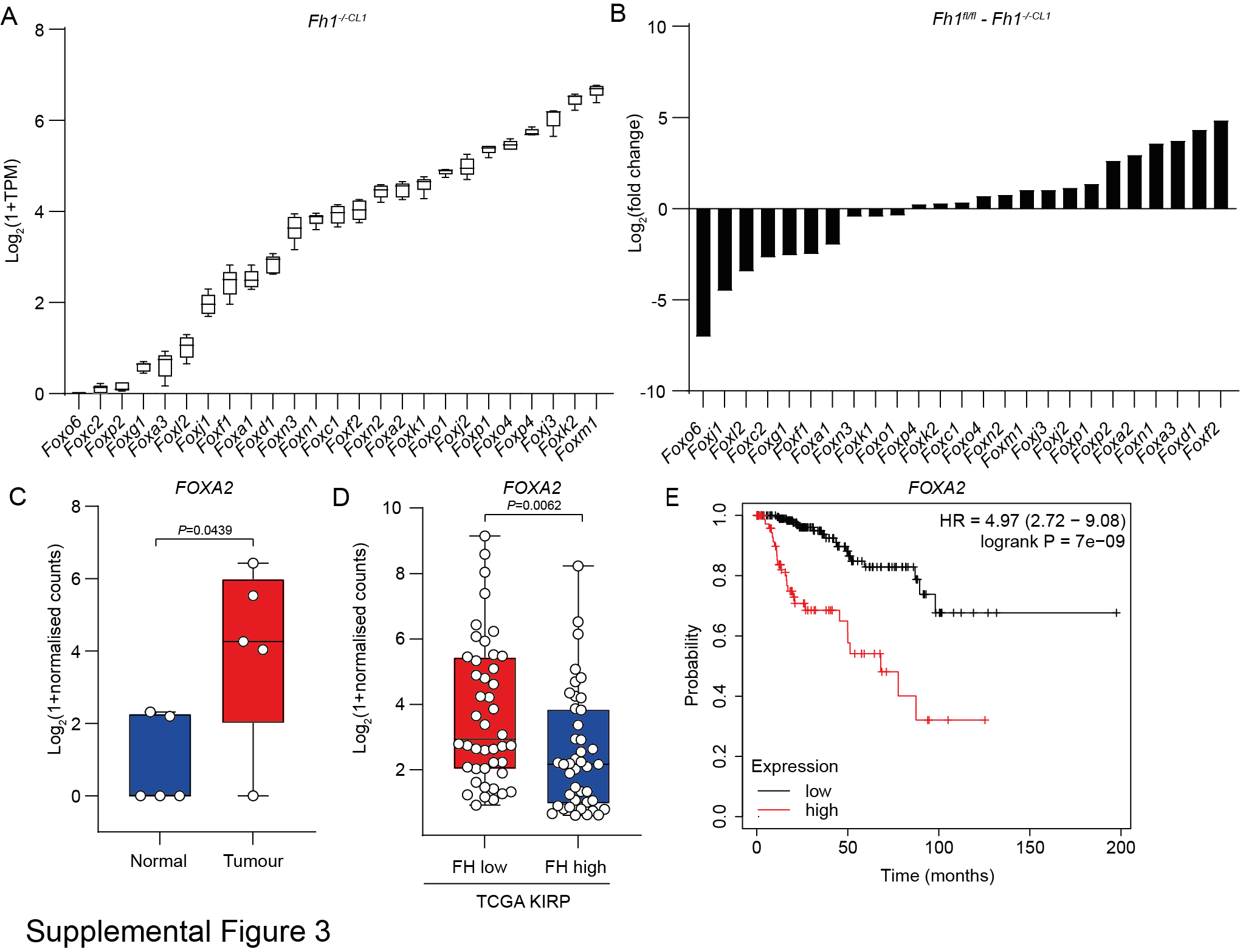

### Supplementary Figure 4

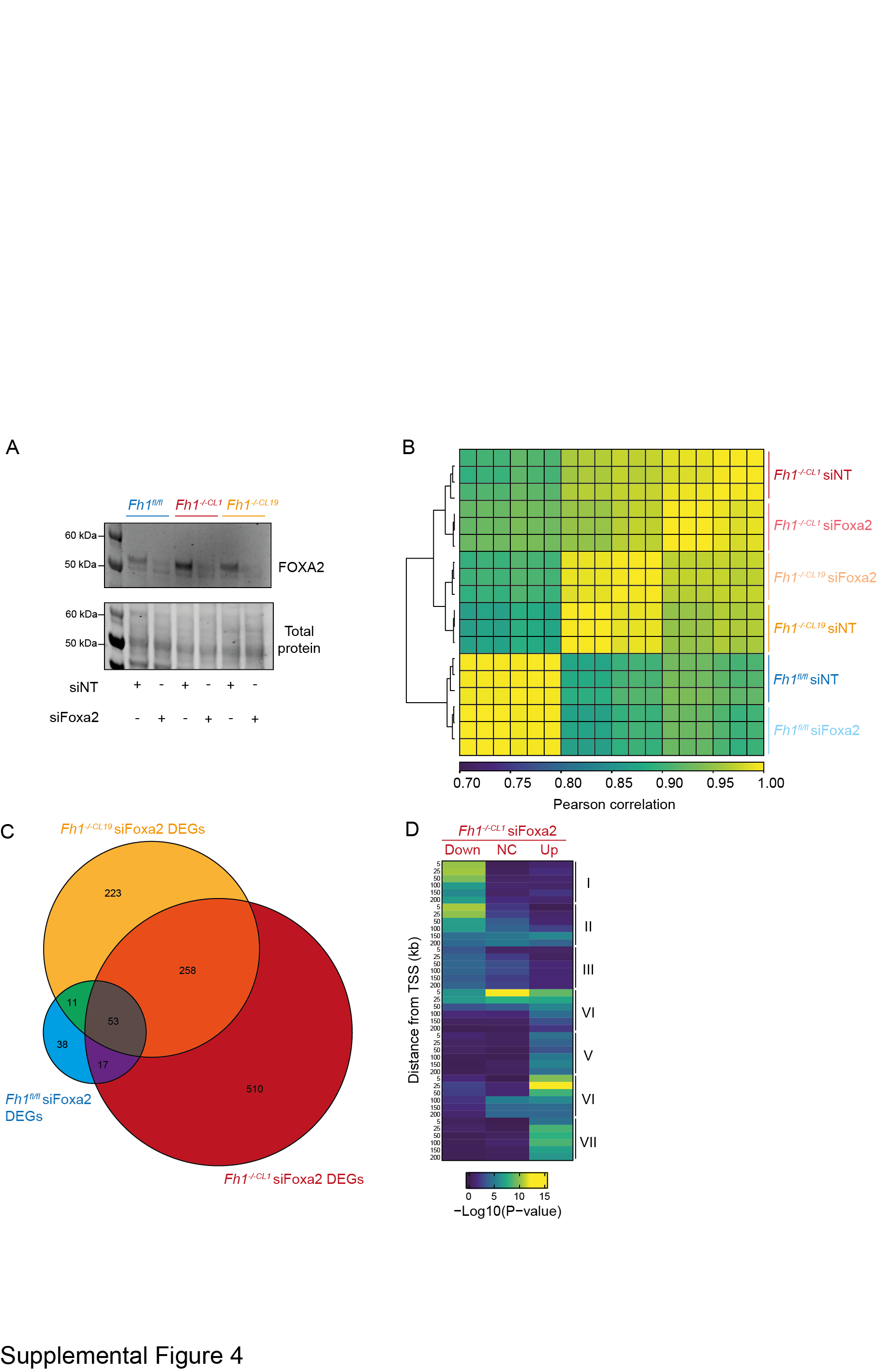

### Supplementary Figure 5

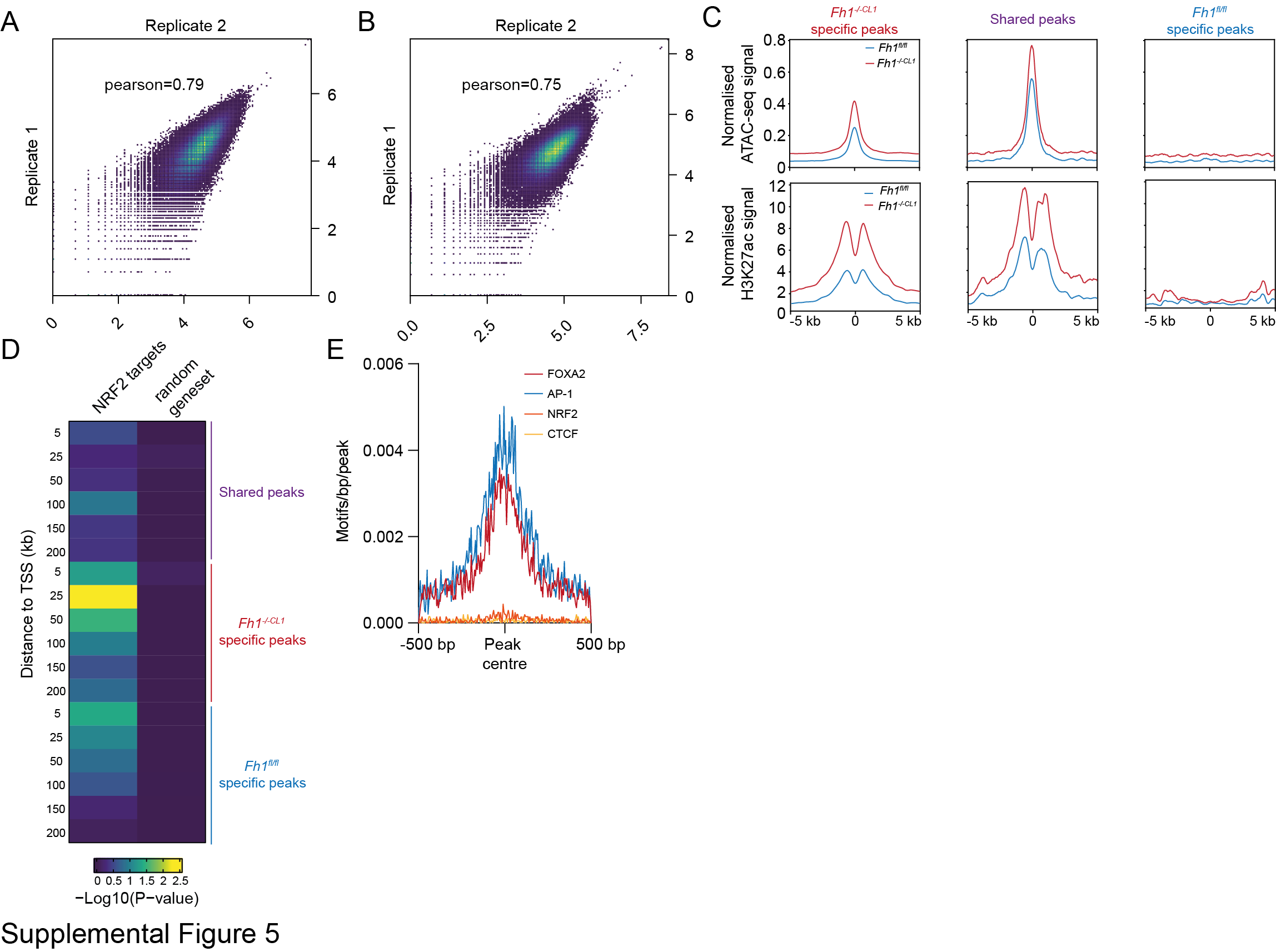

### Supplementary Figure 6

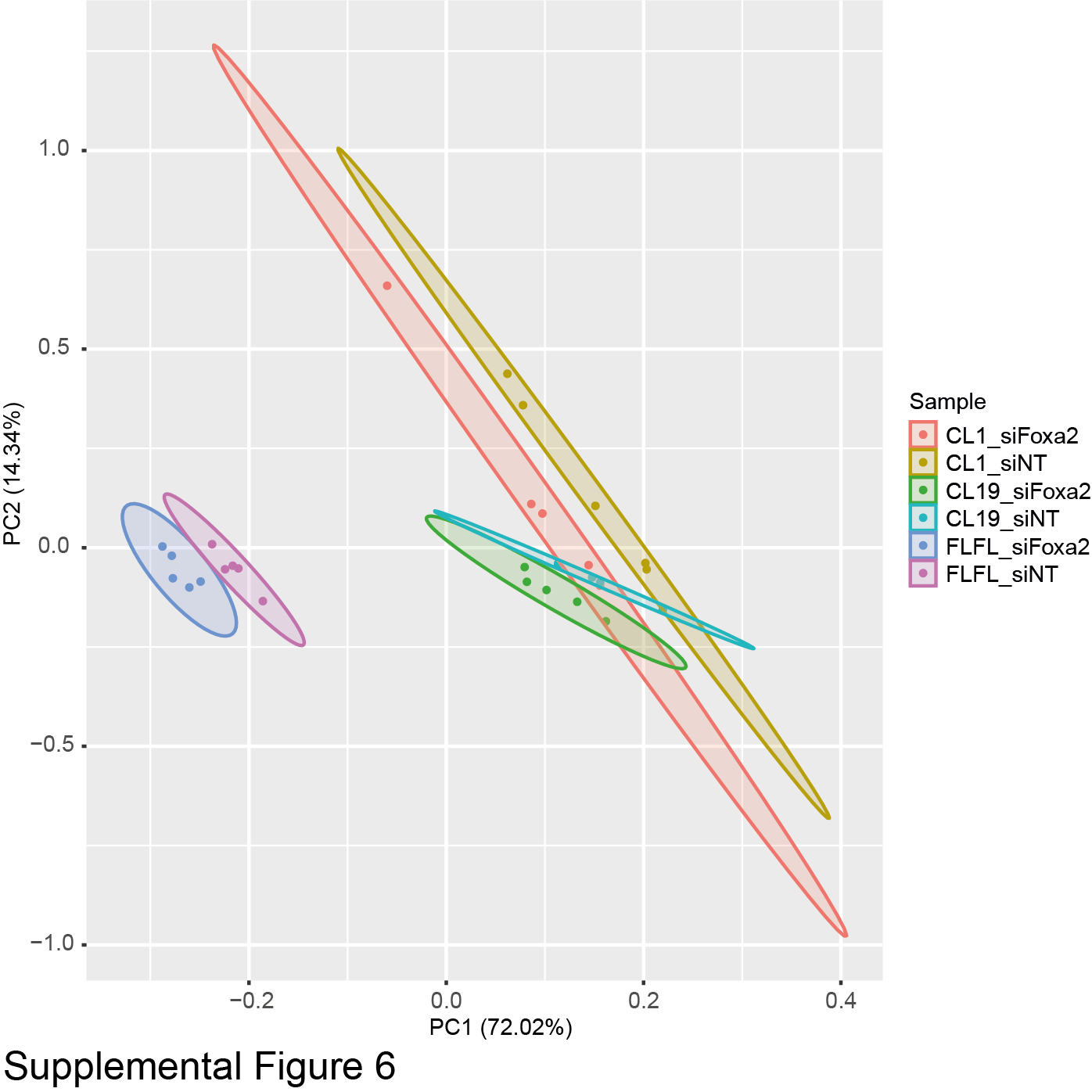
